# Genome-wide identification of novel genes involved in *Corynebacteriales* cell envelope biogenesis using *Corynebacterium glutamicum* as a model

**DOI:** 10.1101/2020.09.29.318030

**Authors:** Célia de Sousa-d’Auria, Florence Constantinesco-Becker, Patricia Constant, Maryelle Tropis, Christine Houssin

**Affiliations:** Université Paris-Saclay, CEA, CNRS, Institute for Integrative Biology of the Cell (I2BC), Gif-sur-Yvette, France; Département Tuberculose & Biologie des Infections, Institut de Pharmacologie et de Biologie Structurale IPBS, UMR5089, Université de Toulouse, CNRS, UPS, Toulouse, France

## Abstract

*Corynebacteriales* are *Actinobacteria* that possess an atypical didermic cell envelope. One of the principal features of this cell envelope is the presence of a large complex made up of peptidoglycan, arabinogalactan and mycolic acids. This covalent complex constitutes the backbone of the cell wall and supports an outer membrane, called mycomembrane in reference to the mycolic acids that are its major component. The biosynthesis of the cell envelope of *Corynebacteriales* has been extensively studied, in particular because it is crucial for the survival of important pathogens such as *Mycobacterium tuberculosis* and is therefore a key target for anti-tuberculosis drugs. In this study, we explore the biogenesis of the cell envelope of *Corynebacterium glutamicum*, a non-pathogenic *Corynebacteriales*, which can tolerate dramatic modifications of its cell envelope as important as the loss of its mycomembrane. For this purpose, we used a genetic approach based on genome-wide transposon mutagenesis. We developed a highly effective immunological test based on the use of anti-arabinogalactan antibodies that allowed us to rapidly identify bacteria exhibiting an altered cell envelope. A very large number (10,073) of insertional mutants were screened by means of this test, and 80 were finally selected, representing 55 different loci. Bioinformatics analyses of these loci showed that approximately 60% corresponded to genes already characterized, 63% of which are known to be directly involved in cell wall processes, and more specifically in the biosynthesis of the mycoloyl-arabinogalactan-peptidoglycan complex. We identified 22 new loci potentially involved in cell envelope biogenesis, 76% of which encode putative cell envelope proteins. A mutant of particular interest was further characterized and revealed a new player in mycolic acid metabolism. Because a large proportion of the genes identified by our study is conserved in *Corynebacteriales*, the library described here provides a new resource of genes whose characterization could lead to a better understanding of the biosynthesis of the envelope components of these bacteria.

## Introduction

The *Corynebacteriales* order is a group of Gram-positive bacteria widely distributed in nature that includes corynebacteria, mycobacteria, nocardia, rhodococci and other related microorganisms [1]. Some of these bacteria are human pathogens, known to cause severe infectious diseases (*Mycobacterium tuberculosis* or *Mycobacterium leprae*) or opportunistic pathologies (*Mycobacterium abscessus*, *Corynebacterium jeikeium* or some species of *Nocardia*). All these bacteria have in common a cell envelope of unusual composition and architecture [2, 3]. Their cell wall core is made up of a peptidoglycan (PG) covalently bound to arabinogalactan (AG) chains, which in turn are linked to mycolic acids (forming the mycoloyl-arabinogalactan-peptidoglycan or mAGP complex). Mycolic acids (MA) are α-branched, β-hydroxylated fatty acids, exclusively synthesized by *Corynebacteriales*, whose length can reach up to 100 carbon atoms in mycobacteria [4]. The mycolic acid-containing part of the mAGP complex associates with other mycolates containing compounds, essentially trehalose mono or di-mycolates (TMM and TDM respectively), to form the backbone of an outer bilayer named the mycomembrane [5, 6]. This outer membrane, that contains porin-like proteins, is thought to be the functional equivalent of the outer membrane of gram-negative bacteria, although it is more impermeable to most compounds and especially antibiotics [7].

Biosynthesis of compounds specific to the cell wall of *Corynebacteriales*, i.e. MA and AG, has been the subject of numerous studies over several decades primarily because the production of these compounds is essential for mycobacterial survival. Hence, AG and MA biosynthesis is the target of several known antituberculous drugs, *e.g.* ethambutol, isoniazid and ethionamide, as well as several candidate drugs in pre-clinical or clinical development [8]. In comparison, *Corynebacterium glutamicum*, a non-pathogenic bacterium widely used in industrial glutamate production, is much more robust against major disruptions of its cell envelope. For example, *C. glutamicum* can grow in the complete absence of MA [9] or with an AG devoid of the arabinose domain [10]. This peculiarity has made this species an indispensable model for the study of the biosynthesis of the *Corynebacteriales* cell envelope. Notwithstanding differences in the fine structure of AM and AG within the *Corynebacteriales*, the major steps of their biosynthesis seem to be conserved among the different genera as evidenced by the presence, in their genome, of genes encoding the enzymes involved in these pathways [11]. Although the cytoplasmic part of these biosynthetic pathways is well understood, a number of unknown factors remain to be discovered regarding the distribution, transit and assembling of these compounds within the cell envelope. Random mutagenesis, using transposons, is a popular approach for identifying such factors. In *Corynebacterium,* only two studies, based on the screening of a transposon-insertion library for mutants with an altered envelope phenotype, have been published [12, 13]. In the first, to identify genes involved in MA synthesis, Wang et al. [12] analyzed approximately 400 insertional mutants of *Corynebacterium matruchotii* using their corynomycolic acid content as a screen. They found one mutant of particular interest with a transposon insertion in a gene encoding a membrane protein conserved in the *Corynebacteriales* (Cg1766 in *C. glutamicum*). However, a subsequent characterization of this protein showed that it is actually an α(1,6) mannopyranosyl-transferase (termed MptB) involved in the synthesis of cell envelope lipoglycans [14]. In a very recent study, Lim et al. [13] generated the first high-density library of transposon insertion mutants of *C. glutamicum* and screened their library for an hypersensitivity to the AG synthesis inhibitor ethambutol. Among the 49 loci identified by their screen (named *ste* for sensitive-to-ethambutol), they found genes encoding proteins already known to be involved in envelope biogenesis but also identified a new locus implicated in cytokinesis.

Because the function of the mycomembrane is to serve as a selective permeability barrier, any defect in the synthesis or assembly of any of the outer membrane components will affect its structure and, consequently, will alter its permeability. Such a relationship between mycomembrane permeability and alteration in MA synthesis has already been shown in *C. glutamicum* using a MytA-deficient mutant [15]. Indeed, disruption of *mytA*, one of the six genes encoding mycoloyltransferases present in the *C. glutamicum* genome [16], produces a significant decrease in cell wall bound corynomycolate and TDM contents, together with an increase in TMM. In this context, Puech et al. [15] showed that the diffusion rate of two hydrophilic molecules was significantly greater in the mutant compared to the parental strain, strongly suggesting an increase in cell wall permeability. We took advantage of this observation to develop a screen that allowed us to readily identify mutants with an altered cell envelope permeability. However, rather than using the passive diffusion of a small molecule through the cell wall to the cytoplasm, as is frequently done in this kind of screen, we searched for the possibility that increased mycomembrane permeabilization could lead to excretion of easily monitored cell wall compounds. For this purpose, we used antibodies directed against AG to screen a transposon mutant library of *C. glutamicum* and identified new genes involved in cell envelope biogenesis, one of which is very likely involved in the biosynthesis of mycolic acids.

## Materials and Methods

### Bacterial strains and growth conditions

The bacterial strains used in this study are shown in Table 1. *Corynebacterium* strains were grown in brain heart infusion (BHI) liquid medium with shaking (250 rpm) or in BHI-agar at 30°C. *Escherichia coli* DH5α was grown at 37°C in Luria-Bertani (LB) medium. When necessary, appropriate antibiotics were supplemented as follows: chloramphenicol (Cm) 15 µg/ml; kanamycin (Km) 25 µg/ml; ampicillin (Amp) 100 µg/ml. Electro-transformable *C. glutamicum* cells were obtained as described in Bonamy et al. [17], with cells collected in early exponential phase (OD_600_ = 1.5) and in the presence of Tween 80 (0.1% v/v final concentration) for strain 2262. Electro-transformable cells were resuspended in 1/500 of the initial culture volume and 100 µl of the cells were pulsed in the presence of 20 to 100 ng of DNA for replicative plasmids, or 500 ng to 3 µg for integrative plasmids (MicroPulserTM electroporator Biorad in 2 mm cuvettes (Eurogentec) at 25 µF, 2.5 kV and 200 Ω). The cell suspension was immediately diluted with 1 ml of BHI medium and incubated for 1h (replicative plasmids) or 2 hours (integrative plasmids) at 30°C before plating.

**Table 1:** Bacterial strains and plasmids used in this study.

| Strain or plasmid | Relevant genotype and/or phenotype | Source or Reference |
| --- | --- | --- |
| *E. coli* strains |  |  |
| *DH5α* | F<sup>-</sup> ϕ80*lacZ*ΔM15 Δ(*lacZ*YA-*argF*)U169 *recA1*<br>*endA1 hsdR*17(*r*<sub>K</sub><sup>-</sup>,*m*<sub>K</sub><sup>+</sup>) *phoA supE*44 λ<sup>-</sup> *thi-1*<br>*gyrA*96 *relA1* | Invitrogen |
| *TOP10* | F<sup>-</sup> *mcrA* Δ(*mrr-hsdRMS-mcrBC*) ϕ80*lacZ*ΔM15<br>Δ*lacX*74 *recA1 araD*139 Δ(*ara-leu*)7697 *galU*<br>*galK rpsL*(Str<sup>R</sup>) *endA1 nupG* λ- | Invitrogen |
| ***C. glutamicum* strains** |  |  |
| 2262 | An industrial strain of *C. glutamicum* | [18] |
| RES167 | Restrictionless derivative of ATCC13032 | [19] |
| CGL2005 | Restrictionless derivative of ATCC21086,<br>Rifampicin resistant | [20] |
| CGL2022 | *mytA* disrupted, Km<sup>R</sup>, derivative of CGL2005 | [15] |
| CGL2029 | Δ*mytA*-Δ*mytB*::Km double mutant, Km<sup>R</sup>,<br>derivative of CGL2005 strain | [21] |
| Δ*cg-pks* | Δ*cg-pks*::Km, Km<sup>R</sup>, derivative of RES167 | [9] |
| Δ*accD3* | Δ*accD3*::Km, Km<sup>R</sup>, derivative of RES167<br>(originally the gene was named *accD4*) | [22] |
| Δ*cg1246* | Δ*cg1246* derivative of RES167 | This work |
| Δ*cg1247* | Δ*cg1247* derivative of RES167 | This work |
| **Plasmids** |  |  |
| pCR<sup>®</sup>2.1-TOPO<sup>®</sup> | *E. coli* cloning vector with *f1* and pUC origins,<br>*lacZα*, Km<sup>R</sup>, Amp<sup>R</sup> | Invitrogen |
| pCGL0040 | pBluescript II SK(+) containing the *cat* gene of Tn9 and the Tn5531 transposon | [23] |
| pK18MobSac | Km<sup>r</sup> *sacB* RP4 *oriT* ColE1 *ori* | [24] |
| pK18MobSac-<br>*Δcg1246* | pK18MobSac containing the upstream and downstream regions of the *cg1246* ORF, construct for *cg1246* deletion | This work |
| pCGL482 | Shuttle vector *E coli*/*C. glutamicum*, Cm<sup>R</sup> | [25] |
| pCGL2420 | Derivative of pCGL482 containing the *cg1246* gene under its own promoter (304 bp upstream the ATG of *cg1247*), Cm<sup>R</sup> | This work |

### DNA manipulations

Plasmid DNA was extracted from *E. coli* using a Wizard® Plus SV Minipreps DNA Purification System (Promega). *C. glutamicum* chromosomal DNA was extracted as described by Ausubel et al. [26]. Oligonucleotide primers were synthesized by Eurogentec. PCR experiments were performed with a 2720 thermocycler (Applied Biosystems) using GoTaq® (Promega) or Phusion^TM^ High Fidelity (Thermo scientific) DNA polymerases. All DNA purifications were performed using a Roche High Pure PCR product purification kit or a QIAquick Gel Extraction Kit (Qiagen). Standard procedures for DNA digestion and ligation were used in conditions recommended by the enzyme manufacturer (Promega or Fermentas). All DNA sequencing was carried out by Beckman Coulter or Eurofins Genomics.

### AG preparation and antiserum production

AG was purified according to [27]. Antiserum against AG was produced in rabbits by CovaLab.

### Mutant generation and immunological screening

Plasmid pCGL0040 (a non-replicative delivery vector containing Tn*5531* [23]) was used to transform fresh electrocompetent *Corynebacterium* 2262 cells [17] and Km resistant transformants were selected on BHI-agar plates supplemented with Km. Subsequently, 10,073 mutants were picked from the original plates and transferred both on BHI-Km-agar plates (12 x 12 cm x 14 mm) entirely covered with a nitrocellulose sheet (Protran® BA-85, Schleicher and Schuell) and to 96-well microtiter plates containing BHI-Km. To grow bacteria, microtiter plates were incubated at 30°C with shaking overnight before sterile glycerol was added (26.5% v/v final concentration). Plates were stored at - 80°C until further use. BHI-Km-agar plates covered with nitrocellulose membranes were also placed at 30°C. Generally, the nitrocellulose sheet was recovered after 16 hours of culture. Nevertheless, when slow-growing colonies were detected, the corresponding mutants were transferred on a new nitrocellulose/BHI plate and allowed to grow for a longer period (24 to 30 h). In order to obtain a replicate of the agar plate, immediately after removal of the nitrocellulose membrane on which the colonies have grown, a new membrane was placed on the plate by gently pressing it to allow a total contact with the agar. After 1 hour of contact at room temperature, the membrane was recovered. All nitrocellulose membranes were treated in the same way. They were washed 3 times for 5 min with phosphate-buffered saline containing 0.05 % (v/v) Tween 20 (PBST), in particular, to completely remove bacteria from the surface of the culture membranes. Membranes were then incubated overnight in blocking buffer (PBST, 5% skim milk powder at 4°C), followed by 3 washes in PBST and a 2 h incubation with primary antibodies against AG (rabbit serum diluted 1:2,000 in PBST). Membranes were then washed 3 times for 5 min in PBST and incubated 1 h with alkaline phosphatase-conjugated secondary antibody (Anti-Rabbit IgG (Fc), AP Conjugate antibody, Promega, 1:7,000 in PBST). After 2 subsequent 5 min washes in PBST, membranes were incubated in a 0.165 mg/ml BCIP (5-bromo-4-chloro-3-indolyl phosphate, Promega)/ 0.33 mg/ml NBT (p-nitroblue tetrazolium chloride, Promega) containing solution (100 mM Tris pH 9.5, 100 mM NaCl, 5 mM MgCl_2_). The reaction development (the appearance of a halo around the colony mark on a culture membrane or of a signal on the corresponding replicate membrane, see above) was followed by comparison to the signal produced by control colonies present on the same culture membrane (the WT strain *C. glutamicum* 2262 and mutants Cg-Pks^-^ and MytA^-^ derived from RES167 and CGL2005, respectively, or from *C. glutamicum* 2262 *i.e.* mutants 1928 and 308). The reaction was stopped by rinsing membranes in deionized water.

At the end of this first screening round, the positive mutants (350) were selected and subjected to the same analysis a second time.

### Identification of disrupted genes containing transposon insertions

Identification of the flanking regions adjacent to the transposon insertions was carried out by inverse PCR [28] or arbitrary-primed PCR [29]. Primers used for these PCR reactions are provided in S1 Table. For inverse PCR, we used the protocol described in Green and Sambrook [28]. Briefly, chromosomal DNAs extracted from the different mutants were digested with either *Sal*I or *EcoR*I restriction endonucleases and then self-ligated with T4 ligase. PCR were performed using primer pairs Isb01/CdsX or Isb04/CdsVIII with the circular DNAs from *EcoR*I or *Sal*I digestions, respectively. Depending on the quantity and purity obtained, PCR products were either purified on columns or purified from agarose gels and, in most cases, sub-cloned into plasmid pCR®2.1-TOPO® using the TOPO TA cloning kit (Invitrogen) prior to sequencing. PCR products were sequenced using primers Isb04 or Isb01 or Rev and F-20. For arbitrary-primed PCR, the first PCR round was performed with approximately 100 ng of genomic DNA as template, using the Phusion High Fidelity DNA polymerase. We used a primer specific for the transposon (Isb01) and a first arbitrary primer that we designed after a search for pentameric sequences present at least 6,000 times in the genome of ATCC13032 strain but absent from the transposon (ARB4020). PCR was performed as follows, with primers at a final concentration of 0.5 µM: 5 min 98°C, 6 cycles (10 s 98°C, 30 s 30°C, 1 min 30 s 72°C), 30 cycles (30 s 98°C, 30 s 45°C, 2 min 72°C) and finally 72°C for 4 min. The PCR products obtained from this first PCR were purified, and one tenth served as template for a second-round reaction. We used a second arbitrary primer (ARBq) that paired with the 5’ end of ARB4020 and a primer that pairs with the transposon downstream of the Isb01 sequence (Isb012). Second-round PCR was performed as follows, with primers at a final concentration of 0.2 µM: 1 min 98°C, 30 cycles (30 s 98°C, 30 s 55°C, 2 min 72°C) and finally 72°C for 4 min. If only one major band was visible on an agarose gel after this second-round PCR, the product was purified and sequenced. However, when several bands of comparable intensity were present on an agarose gel, the different products were purified from the gel and submitted to a third round of PCR in conditions identical to that of the second round. PCR products were sequenced using primer Isb013.

### Bioinformatic analyses

Sequences interrupted by the transposon were aligned using BLASTn online software at the NCBI (National Center for Biotechnology Information; http://www.ncbi.nlm.nih.gov) using *Corynebacterium* genomes as the search set. Best matches were systematically obtained with sequences from SCgG1 and SCgG2 genome strains. However as neither of these two genomes is annotated, transposon insertion locations were established with respect to the sequence of the ATCC13032 genome (NCBI Reference Sequence: NC_006958.1).

Conserved domains, or functional units within proteins, were searched using the Conserved Domain Database (CDD, https://www.ncbi.nlm.nih.gov/Structure/cdd/wrpsb.cgi) at the NCBI [30] and the Integrated Microbial Genomes and Microbiomes web resources (IMG/M: https://img.jgi.doe.gov/m/) [31]. Unknown proteins were classified into general categories using EggNOG (Evolutionary genealogy of genes: Non-supervised Orthologous Groups) [32] (http://eggnog5.embl.de).

Analysis of the *C. glutamicum* transcriptome published by Pfeifer-Sancar et al. [33] was used to predict genes inactivated in operons.

### Construction of plasmids and bacterial strains

The different plasmids used in this study are described in Table 1. In order to delete *cg1246*, we used the strategy described by Schafer et al. [24]. In brief, two DNA fragments overlapping the gene at its 5’ and 3’ extremities were amplified by PCR from *C. glutamicum* total DNA using appropriate primers (1246-del1/1246-del2 and 1246-del3/1246-del4, see S1 Table) and cloned in the non-replicative vector pK18mobSac. The resulting plasmids (pK18mobsacΔ1246) was sequenced and transferred into *C. glutamicum* RES167 by electroporation. Transformants in which the construct was integrated into the chromosome by single crossing-over were selected on BHI plates containing Km. The second crossover event was selected by plating Km^R^ clones on BHI plates containing 10 % sucrose. Km-sensitive and sucrose-resistant colonies were screened by PCR for the correct deletion of the gene using appropriate primers. After verification of PCR products by sequencing, one strain carrying the *cg1246* deletion (Δ1246 strain) was selected for further studies.

A complementation vector encoding Cg1246 (pCGL2420) was constructed using pCGL482 as the cloning vector [25]. Two DNA fragments were amplified by PCR from *C. glutamicum* RES167 chromosomal DNA: the coding sequence of *cg1246* (using the primer pair 1246-RcaI/1246-XhoI) and the promoter region of the operon *cg1247-cg1246* (304 bp upstream the ATG of *cg1247*, using the primer pair p1246-RcaI/p1246BglII) (see S1 Table). The amplicons were ligated together, digested with *Rca*I and *Xho*I and inserted into the *BamH*I/*Xho*I digested pCGL482 to obtain plasmid pCGL2420.

### Biochemical analyses of selected mutants

#### Protein analyses

Cells grown overnight (equivalent of 1 ml bacterial suspension at an OD_650_ = 10) were centrifuged at 16,000 g for 5 min. Then, 800 µl of the supernatant, which contained the secreted proteins, were added to 200 µl of 50% TCA and the mixture incubated for 1 h at 4 °C. The precipitated proteins were collected by centrifugation, the pellet was washed with acetone and solubilized in 50 µl of Laemmli denaturing buffer. The bacterial pellet was incubated in 50 µl of 50 mM Tris-HCl pH 6.8, 2% (w/v) SDS at 100 °C for 3 min and centrifuged at 16,000 g, for 5 min at 4 °C. The supernatant, which contained the cell wall proteins, was recovered. Proteins were separated by SDS-PAGE and gels were stained with Coomassie brillant blue R-250.

#### Lipid analysis

Lipids were extracted from wet cells for 16 h with CH_3_OH/CHCl_3_ (2:1 v/v) at room temperature. The organic phase was evaporated to dryness and lipids were solubilized in CHCl_3_ (typically 100 µl for lipids extracted from 20 ml of exponentially growing cells or 10 ml of cells in stationary phase). Lipids were analysed by Thin Layer Chromatography (TLC) on silica gel-coated plates (G-60, 0.25 mm thickness, Macherey-Nagel) developed with CHCl_3_/CH_3_OH/H_2_O (65:25:4, v/v/v). Detection of all classes of lipids was performed by immersion of the TLC plates in 10% H_2_SO_4_ in ethanol, followed by heating at 110°C; glycolipids were revealed by spraying plates with 0.2% anthrone (w/v) in concentrated H_2_SO_4_, followed by heating at 110°C.

The various classes of extractable lipids were also analysed by TLC after radiolabelling. Briefly, 1 µCi of [1-^14^C]-palmitate (2.22 GBq mmol-1, Perkin Elmer) was added to 10 ml culture medium of exponentially or stationary phase-grown bacteria and further incubated for 1.5 h at 30°C. After centrifugation, the cell pellets were extracted twice with CHCl_3_/CH_3_OH (1:2, v/v, then 2:1, v/v) for 2×24 h. The organic solutions were separated from the delipidated cells by filtration, then pooled and dried. The crude lipid extracts were resuspended in CHCl_3_ at 20 mg/mL and 15 µL were spotted onto a Silica Gel 60 TLC plate run in CHCl_3_/CH_3_OH/H_2_O (65:25:4, v/v/v). Labelled lipids were visualized with a Typhoon phosphorImager (Amersham Biosciences). The relative percentage of radioactivity incorporated in TMM and in TDM was determined using Image Quant software (GE Healthcare).

## Results and Discussion

### Development of an original screen to detect mutants with envelope disruption

To identify new genes involved in *Corynebacterium* cell envelope biogenesis, we developed an effective and rapid test using polyclonal antibodies directed against AG. For this purpose, we took advantage of an observation that we made with well-characterized mutants partly or totally devoid of mycolic acids. Indeed, as shown in Fig 1, these mutants released, to the external medium, molecules that reacted with anti-AG antiserum. This excretion was clearly visible as a halo around a colony when bacteria were spotted and grown on a nitrocellulose membrane on a BHI plate, after immunoblotting with anti-AG antiserum. No signal was visible when the corresponding pre-immune serum was used instead of the anti-AG antiserum (data not shown). Moreover, the size of the excretion halo is commensurate with the importance of the alteration, as can be seen from a comparison of the wild type (WT) strain, the MytA^-^ mutant [16], the double MytA^-^/MytB^-^ mutant [21] and the Cg-Pks^-^ [9] or AccD3^-^ mutants [22] which produce 100%, 60 %, 40 % or no mycolic acids, respectively. The same pattern was also obtained with 2 other mutants defective in AG biosynthesis: AftB^-^ [34] and DprE2^-^ [35] (data not shown). Although we were unable to identify the exact nature of the excreted compound(s) reacting with the anti-AG antibodies (probably because of the small amount present in the external medium), we believe that this information was not essential for mutant screening and that this assay could thus be a very powerful means to identify new genes involved in envelope biogenesis.

**Figure 1:**
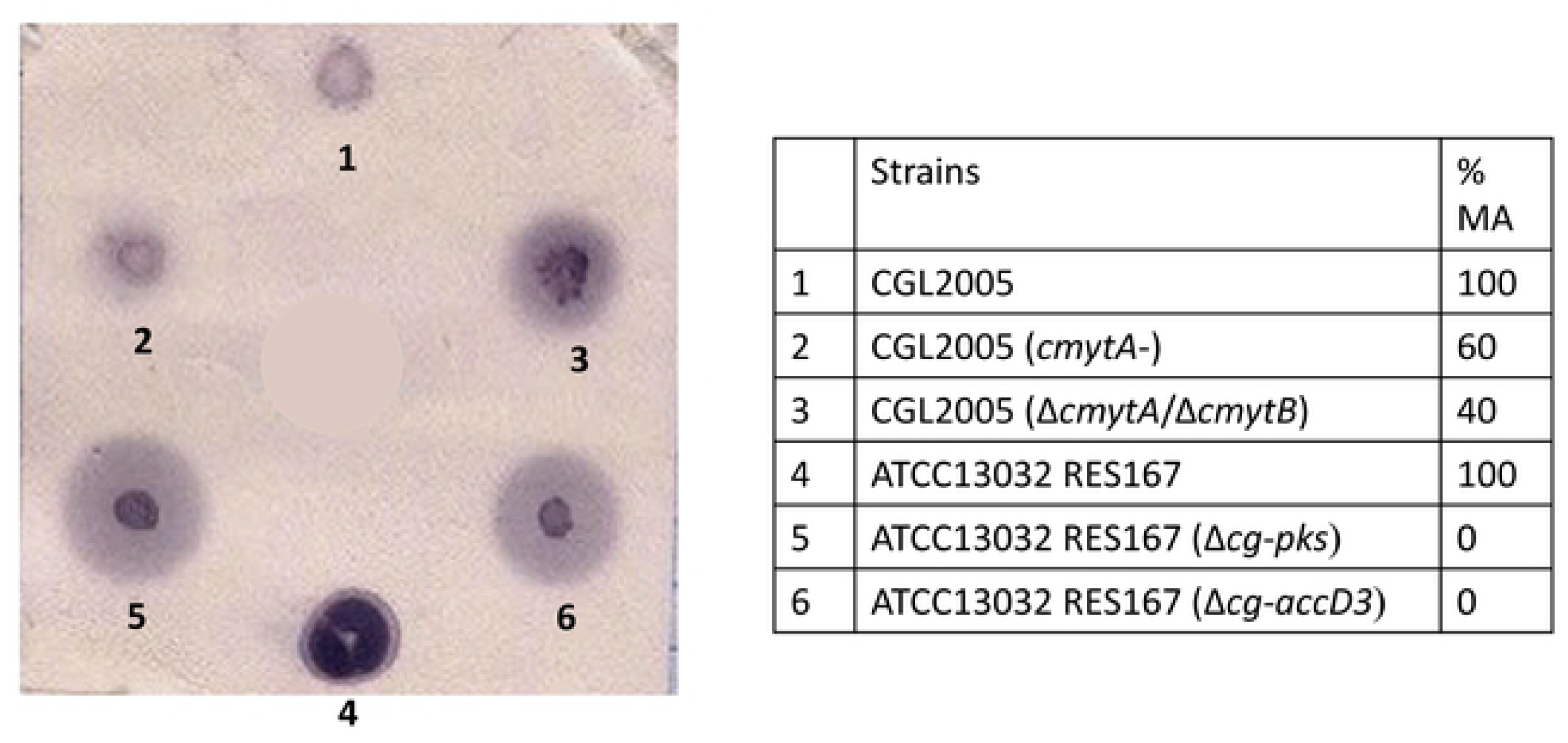
Immunological test for the detection of mutants with an altered cell envelope permeability. Different *C. glutamicum* strains were spotted and cultivated on a nitrocellulose membrane placed on a BHI plate as described in Materials and Methods. The membrane was first treated with primary antibodies against AG and then by a classical western blot procedure (phosphatase alkaline coupled secondary antibodies and NBT/BCIP revelation). Clone outlines are colored in dark purple and are visible for all strains. Mutants are surrounded by a very distinct halo (light purple around the colony), the size of which is proportional to the importance of the cell envelope perturbation. The strains used in this test (numbered 1 to 6 in the blot) are given in the table. The % MA indicates the proportion of MA in the mutants as compared to the corresponding parental strain.

### Construction of a transposon library in *C. glutamicum* and large-scale immuno-screening of mutants with an altered envelope

A transposon mutant library of *C. glutamicum* was generated using an IS*1207*-based transposon, Tn*5531*, cloned into a non-replicative delivery vector [23]. This system was shown to be effective for random mutagenesis in *C. glutamicum* 2262, a strain that does not contain the IS*1207* sequence in contrast to the reference strain ATCC13032 ([23] and unpublished results). We generated 10,073 insertion mutants in this strain that we analysed by means of our immunological test.

The scheme of the library screening is depicted in Fig 2. Because differences in growth kinetics of mutants and in plate humidity could generate variability in the diffusion rate of the antigenic molecules from the nitrocellulose to the agar, we tested each mutant for both the presence of a halo around the colony and/or the presence of a positive signal on the plate footprint (S1 Fig). To ensure reliable immunological responses, two rounds of screening were performed. In this way, we were able to select 133 mutants that unambiguously excrete into the medium a compound recognized by anti-AG antiserum. In order to refine these data, we searched for characteristics that are commonly observed in cell wall mutants and that could be present in these pre-selected mutants. These mainly concerned (i) growth and phenotype differences and (ii) differences in cell wall and secreted protein profiles as compared to the parental strain. For the first parameter, we analysed colony phenotypes on agar plates, the rate of growth on solid and liquid media, the tendency to aggregate in liquid culture and the presence of cell shape or division defects visible by optical microscopy. The second parameter was based on observations previously made that an alteration of cell-wall architecture makes the strain more sensitive to SDS treatment, leading to the extraction of a greater number of cell envelope proteins [21]. Cell envelope alteration also often produces a leakage of cell wall proteins into the culture medium [16]. We then sorted the mutants by assigning them a score as follows: we rated 1 or 2 the immunological signal produced by a mutant according to the size of its diffusion halo (see S1 Fig). We assigned 1 point to each mutant exhibiting at least one of the phenotypic or growth differences listed above and 1 point to each mutant with significant differences in cell wall/secreted protein profiles (see S2 Fig for examples). After summing these three scores, only mutants with a total score ≥ 2 were retained, reducing their number from 133 to 80 (i.e. 0.8% of all the 10,073 mutants analysed by our immunological screening). Details of the scores attributed to each of the mutants finally selected are provided in S2 Table.

**Figure 2:**
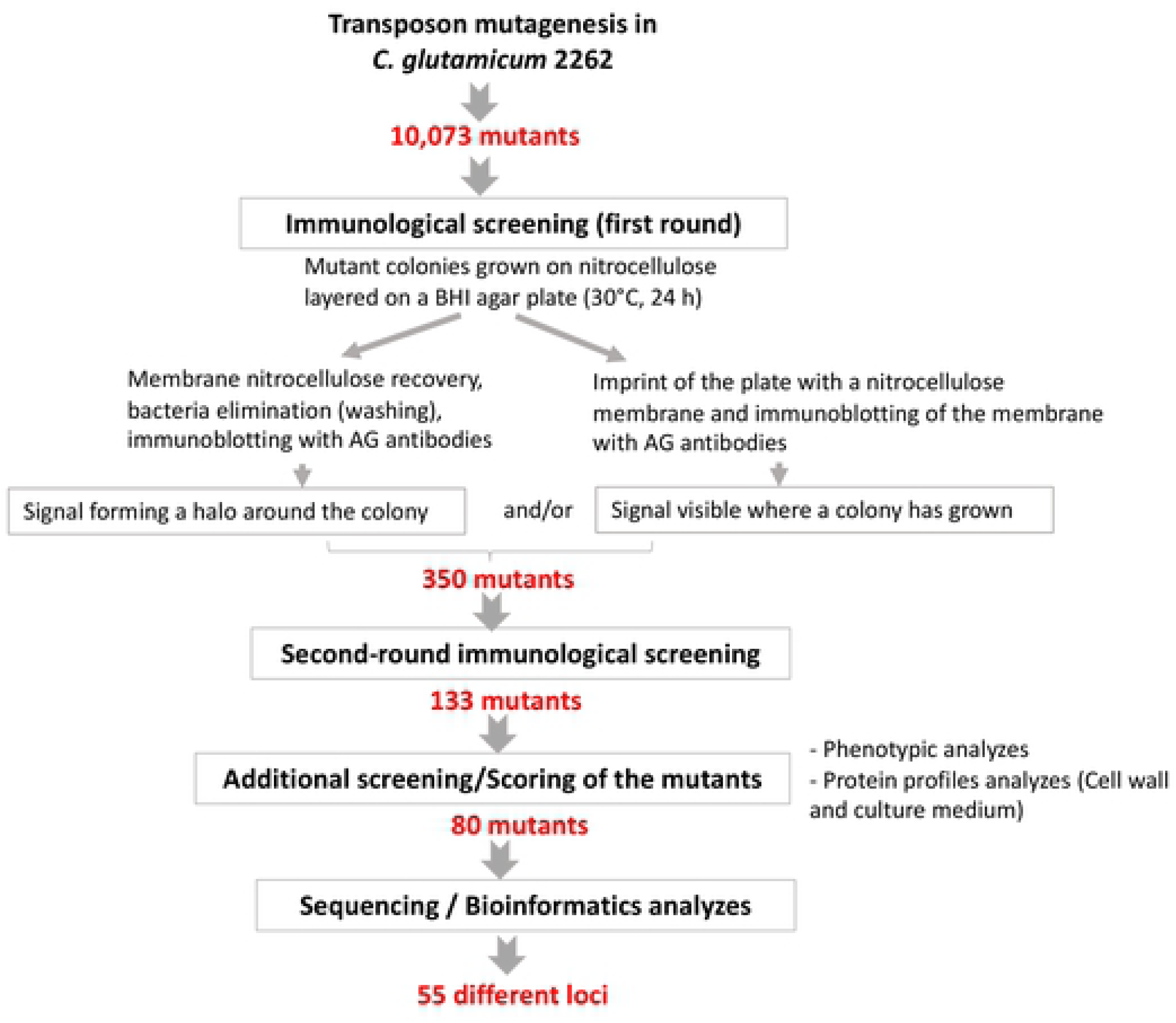
Flow chart of the screening procedure used in this study. See Materials and Methods for details.

### Analyses of Tn insertions

The precise insertion site of the transposon in the genome of each of these 80 mutants was then determined by PCR analyses (inverse or arbitrary PCR) and DNA sequencing. Because the sequence of the *C. glutamicum* 2262 strain is not available, a nucleotide BLAST analysis was performed for each of the amplicons against all *C. glutamicum* genomes available in the National Center for Biotechnology Information (NCBI) database. In all cases, similar DNA sequences were found with the highest identity scores systematically obtained for strains SCgG1 (NC_021351.1) and SCgG2 (NC_021352.1), two industrial glutamate hyper-producing strains. As shown in Fig 3A, the selected mutations were distributed all along the chromosome, but with a number of loci hit several times and a substantial number of insertions in a region known to be involved in envelope biosynthesis (Fig 3A and 3B, see below). Of the 80 sequences interrupted by the transposon, 79 could also be unambiguously mapped on the ATCC13032 strain chromosome sequence. The only sequence that could not be mapped was identified at the upstream region of the *mytC* gene orthologue in SCgG1 and SCgG2. Because the ATCC13032 strain is the most documented of the *C. glutamicum* strains, we chose to refer to it, and in particular to the NCBI Reference Sequence: NC_006958.1 [36]. In this context, and in the absence of genome annotation for *C. glutamicum* 2262, we annotated genes in this strain by the locus tag identifier and the gene symbol of the orthologous locus found in ATCC13032 preceded by ort*-* for the locus tag.

**Figure 3:**
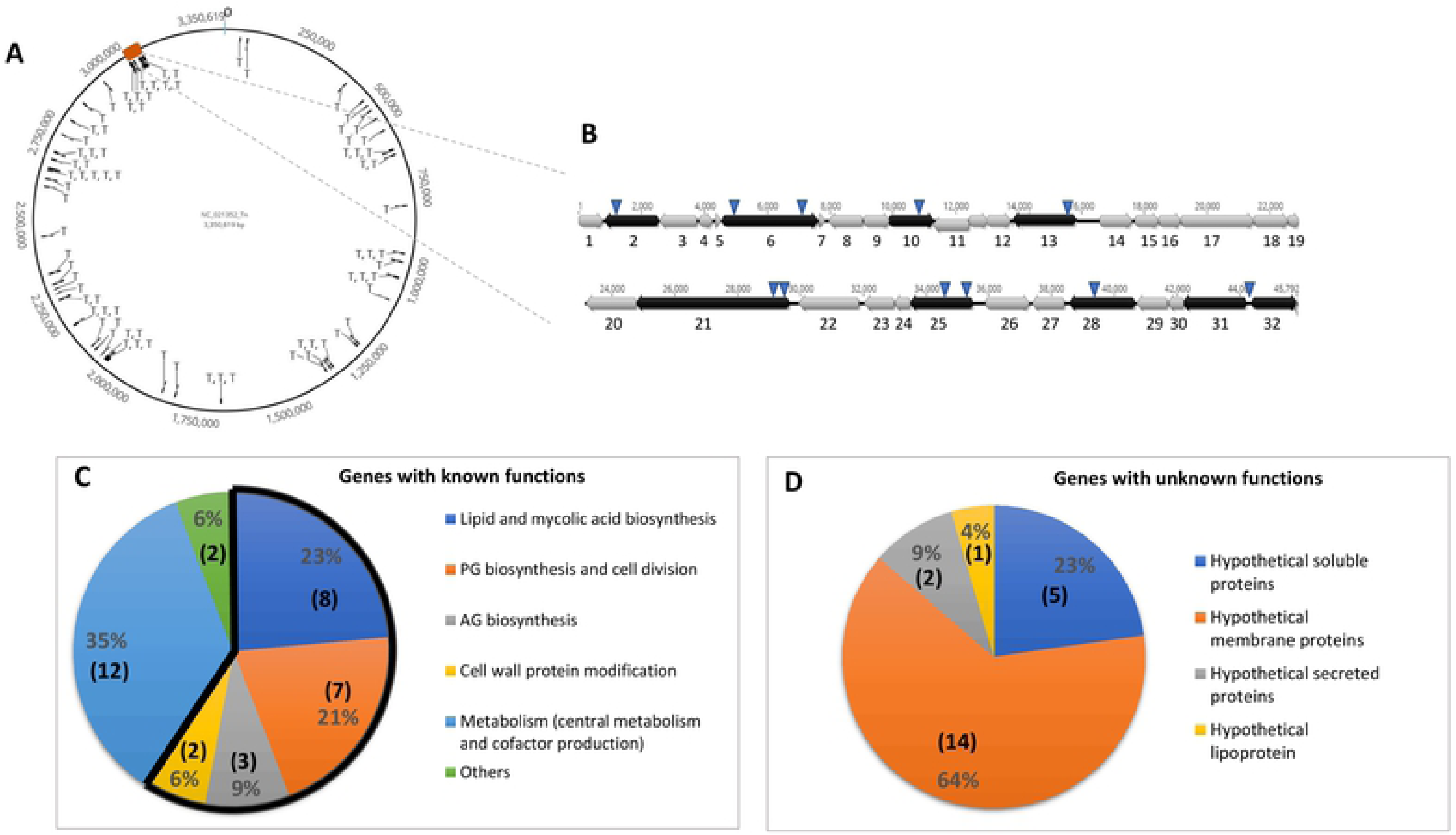
Overview of the mutant analyses. (A) Location of transposon insertions (indicated by T) corresponding to the 80 selected mutants, mapping to the genome of *C. glutamicum* SCgG2. The numbers outside the circle represent the base pairs (from 0 to 3,350,619). The red rectangle corresponds to the large cluster of genes known to be involved in cell envelope biogenesis, which is detailed in (B). (B) Schematic representation of the cell wall biosynthetic gene cluster of *C. glutamicum* ATCC13032. This large cluster includes many genes involved in mycolic acid and AG biosynthesis. The cluster is given from *cg3156* to *cg3192* (about 45.8 kb) but its limits are not precisely known. The distances on the chromosome are indicated. Gene orientation is indicated by arrows; genes that were insertionally-inactivated are in black; the locations of transposon insertions are indicated by inverted triangles. For better readability, the genes are numbered from 1 to 32. Details of their annotations (locus tag, protein name and function) are provided below. 1: *cg3156* (HsecP), 2: *cg3157* (HsecP), 3: *cg3158* (NagA2: putative β-N-acetylglucosaminidase), 4: *cg3159* (UspA), 5: *cg3160* (HsecP), 6: *cg3161* (AftD, arabinosyltransferase, AG biosynthesis), 7: *cg3162* (HP), 8: *cg3163* (TmaT, TMCM mycolylacetyltransferase), 9: *cg3164* (HMP), 10: *cg3165* (HMP), 11: *cg3166* (HP, putative glycosyltransferase), 12: *cg3167* (HP), 13: *cg3168* (MtrP, methyltransferase), 14: *cg3169* (PhosphoenolPyruvate Carboxykinase), 15: *cg3170* (Tellurite resistance protein or related permease), 16: *cg3172* (tRNA (guanine-N(7)-)-methyltransferase), 17: *cg3173* (HP), 18: *cg3174* (CmpL1 mycolic acid transporter), 19: *cg3175* (HMP), 20: *cg3176* (HP), 20: *cg3177* (AccD3, subunit of Acyl-CoA carboxylase complex, mycolic acid biosynthesis), 21: *cg3178* (Cg-Pks, mycolic acid condensase), 22: *cg3179* (Cg-FadD2, fatty acyl-AMP ligase, mycolic acid biosynthesis), 23: *cg3180* (Elrf, envelope lipid composition regulator), 24: *cg3181* (HSecP), 25: *cg3182* (MytA, mycoloyltransferase), 26: *cg3185* (Pcons, HP), 27: *cg3186* (MytB, mycoloyltransferase), 28: *cg3187* (AftB, AG biosynthesis), 29: *cg3189* (UbiA, decaprenylphosphoryl-D-arabinose (DPA) biosynthesis), 30: *cg3190* (5’-phosphoribosyl-monophospho-decaprenol phosphatase, DPA biosynthesis), 31: *cg3191* (Glft2, galactosyltransferase, AG biosynthesis), 32: *cg3192* (HMP). (C) Pie chart representing the distribution, by functional categories, of the proteins of known function identified by our screening procedure. The part of the circle outlined in black represents the categories that are directly related to the biogenesis of the cell envelope. Numbers in parenthesis represent the number of genes in the corresponding category. (D) Pie chart representing the distribution, by putative localizations, of the proteins of unknown or poorly characterized functions identified by our screen. Numbers in parenthesis represent the number of genes in the corresponding category.

In 70 % of the mutants, the transposon insertion was found to occur inside an open reading frame (ORF). However, in the remaining 30%, transposon insertion site was found in a non-coding region at the 5’-end of an ORF (between 10 base pairs (bp) and up to 200 bp upstream of the start codon, depending on the mutants), that we assumed to be the promoter region (noted pr-), the interruption of which may, more or less dramatically, affect downstream ORF expression.

Of these 80 mutants, only 55 corresponded to different disrupted loci. Indeed, in 15 cases the same ORF and/or promoter region was found to be disrupted by the insertion in 2 (7 cases), 3 (7 cases) or 5 times (1 case). Nevertheless, in 3 cases, several insertions were in the exact same position (3 in pr-*fasI*, 2 in pr-*lgt* and 2 in *steA*, see S2 Table). We cannot exclude that identical insertions came from cross-contaminations although this seems unlikely because the different strains did not originate from the same plates and that, at least for the *fasI* and *steA* mutants, the transposon (and consequently the *aphIII* gene) was not inserted in the same direction in all the mutants. The difference in the orientation of the *aphIII* gene at the 5’ end of the *fasI* gene probably lead to different polar effects that could explain the variations observed in the scores obtained for the three mutants with the same transposon insertion point.

It has been shown that only one-third of the approximately 3000 protein-coding genes of the ATCC13032 strain are transcribed monocistronically while the remaining two-thirds are part of operons [33]. If we assume that the transcriptional pattern of *C. glutamicum* 2262 is similar to that of the ATCC13032 strain, then, 35 of the interrupted loci lie within operons, which represent 64% of the total number of impacted loci, as would be expected if the transposon was randomly inserted into the genome.

All information concerning the mutants obtained in our library (score, transposon insertion sites, locus tags, prediction of a gene in an operon, characteristic of gene products) are given in S2 Table.

### Immuno-screening with anti-AG antibodies is effective to identify genes involved in cell wall biogenesis of *Corynebacteriales*

Bioinformatic analyses of DNA sequences interrupted by the transposon showed that 34 loci (51 different mutants) correspond to ORFs or promoter regions of previously characterized genes (Fig 3C). These loci are shown in Table 2. Among them, 20 are directly involved in cell wall processes. Two genes encode proteins that synthetize essential molecules for cell envelope building: (i) fatty acids, the precursors of phospholipids and mycolic acids (5 hits) and (ii) decaprenyl-pyrophosphate, the lipid carrier of many precursors of cell wall compounds (insertion in the promoter region of *uppS1*). Thirteen interrupted loci were found to be directly involved in mAGP complex biosynthesis: *cg*-*pks*, pr-*pptT, cmrA*, pr-*dtsR2*, *mytA, mtrP* (mycolic acid biosynthesis), *aftB*, *aftD,* pr*-glfT2* (AG biosynthesis), *ponA*, *ftsI*, *alr*, *ltsA* (PG biosynthesis). Three transposon insertions affected genes involved in cell division (*fhaA*, *ftsK* and *mraW* which, with *ftsI*, belongs to the *dcw* cluster). Two loci encoding enzymes that modify envelope proteins post-translationally were also interrupted: *lgt*, and pr*-mytC*, responsible for the transfer of a diacylglyceride or a mycolate onto proteins, respectively. Twelve of the 14 other genes are mainly related to metabolic functions and energy production, most of which may have an indirect influence on the biosynthesis of envelope compounds. This is most probably the case for genes encoding enzymes of central metabolism (*lpd*, *mqo*, *deoC* and *zwf*) or encoding proteins involved in the assembly of the cytochrome *bc1–aa3* supercomplex (*ctiP* and pr-*ctaD* and *surf1*), the inactivation or under-expression of which will certainly modify respiratory chain activity and consequently carbon fluxes. Two genes (*otsA* and *otsB*), belonging to one of the three different trehalose synthesis pathways present in *C. glutamicum*, were also interrupted by the transposon. Because trehalose is the main acceptor of mycolates in the cell envelope [37], it is not very surprising to obtain such mutants by our screening. We found 3 independent insertions in *pdxR*, a gene encoding a positive transcriptional regulator of the pyridoxal 5’-phosphate synthase genes. Many enzymes use pyridoxal 5’-phosphate (PLP) as a cofactor, primarily thus involved in the biosynthesis of amino acids and their derivatives. For example, *meso*-diaminopimelate (*m*-DAP), an essential amino-acid in PG, is synthetized by two pathways, one of which (the succinyl pathway) uses DapC (the succinyl-diaminopimelate transaminase), a PLP-dependent enzyme, and it has been shown that mutations in this pathway led to loss of cell wall integrity [38]. We also found an insertion in *bioM*, which is part of the *bioYMN* operon encoding the biotin transport system. *C. glutamicum* is a biotin auxotrophic bacterium, and must import the cofactor from its environment by the ATP-dependent BioYMN transport system [39]. Biotin is essential for acyl-CoA carboxylases involved in fatty acid and mycolate biosynthesis and it has been shown that biotin limitation can lead to a small decrease in mycolic acid content but also to an important change in their chain length [40].

**Table 2:**
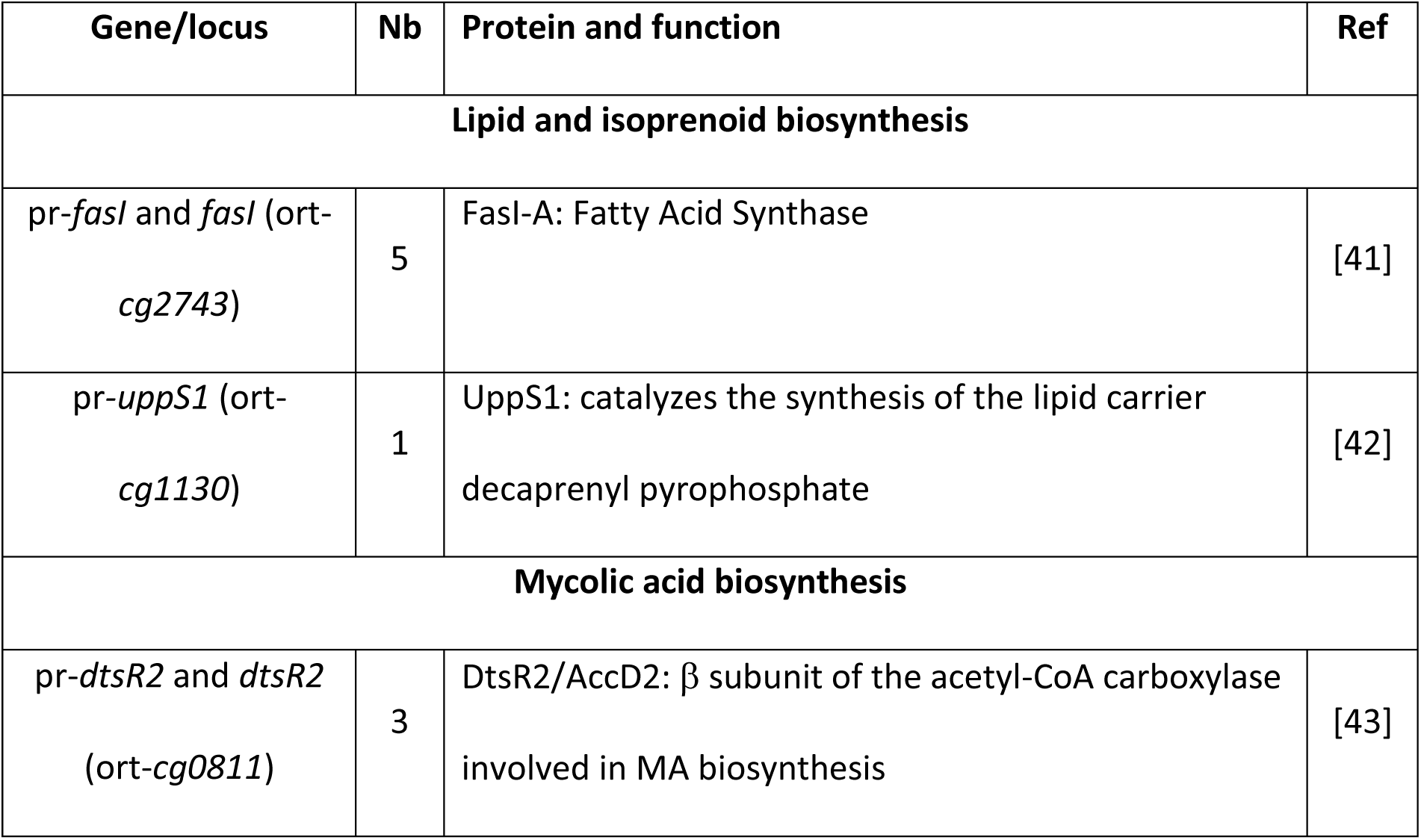
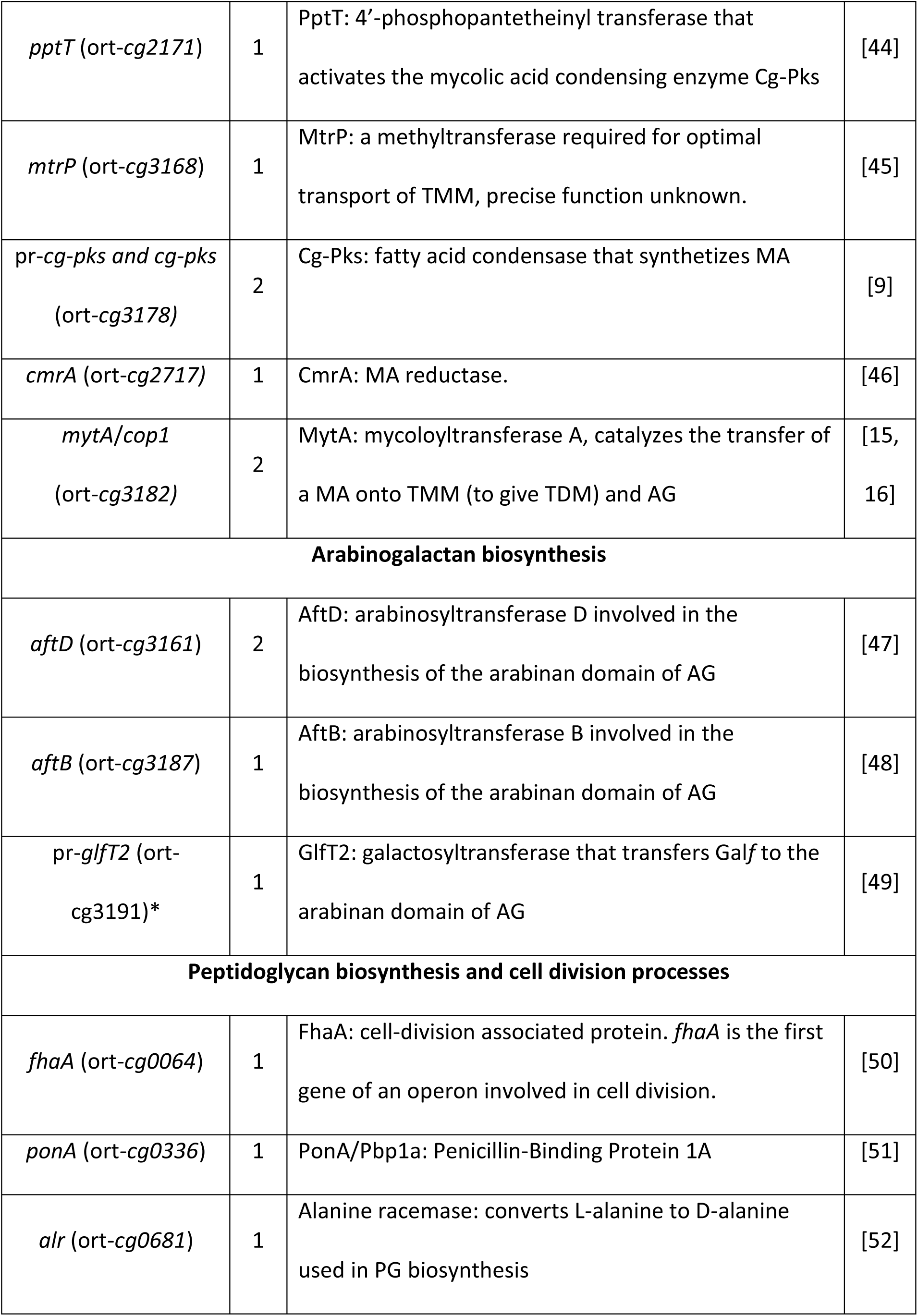
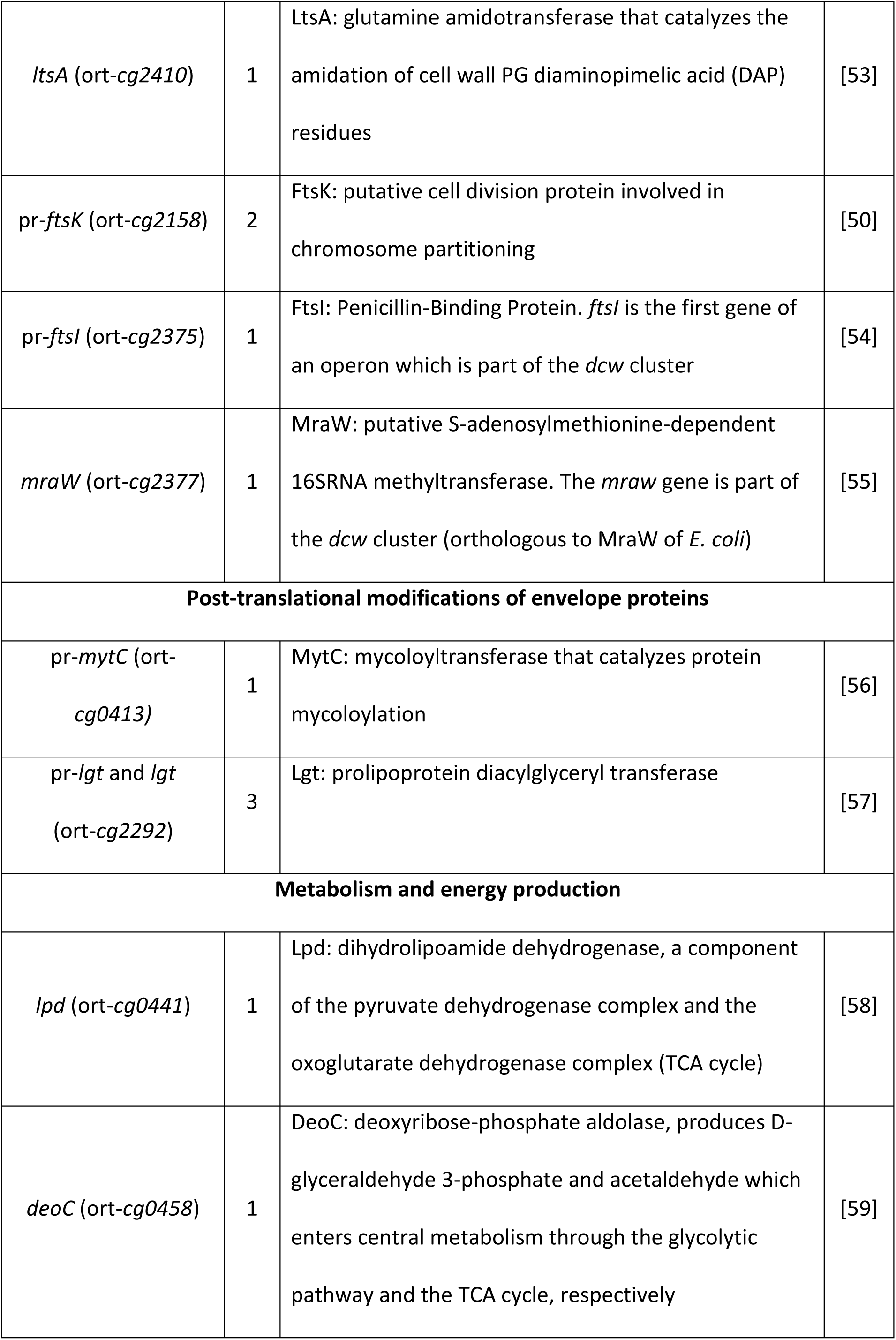
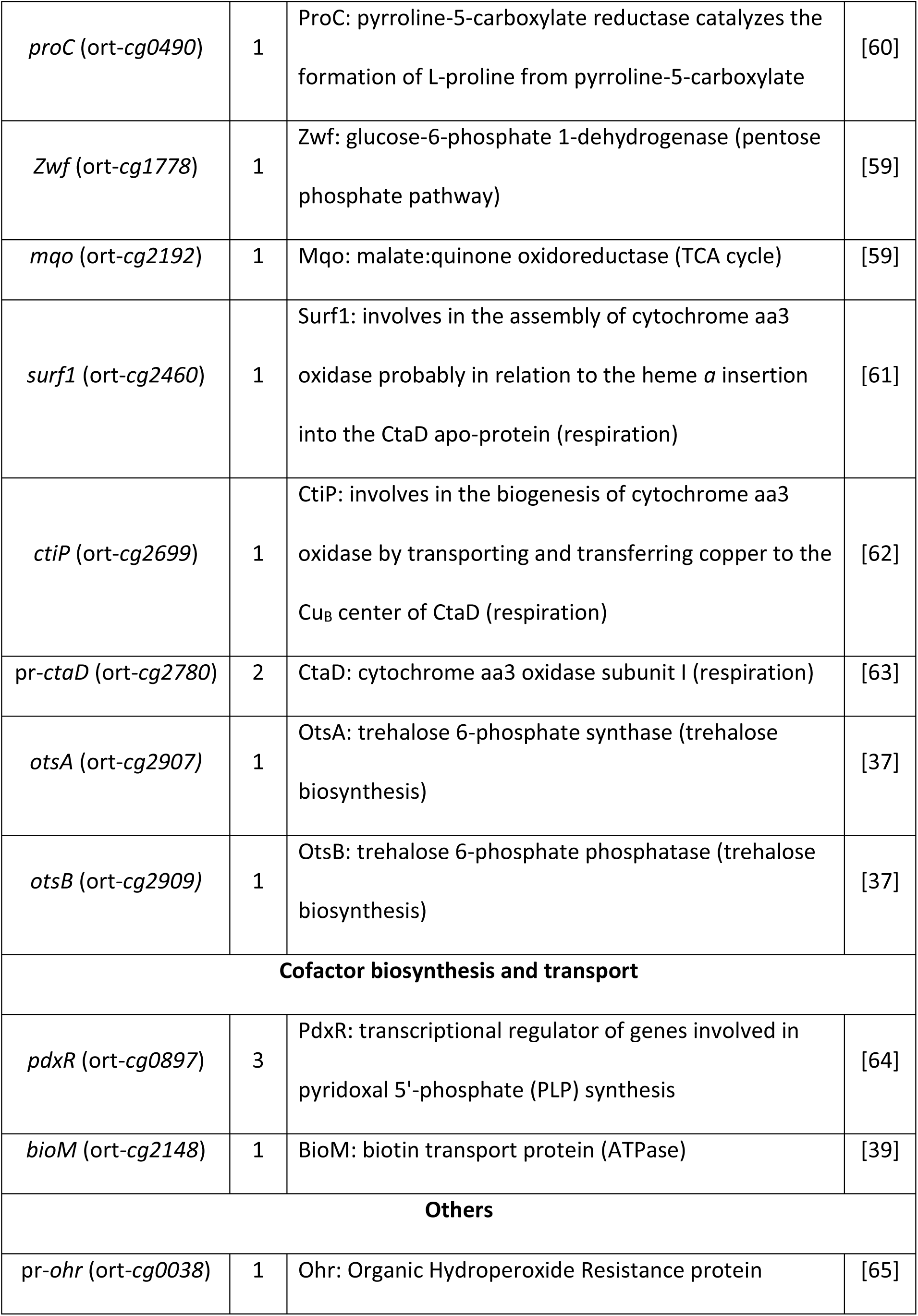
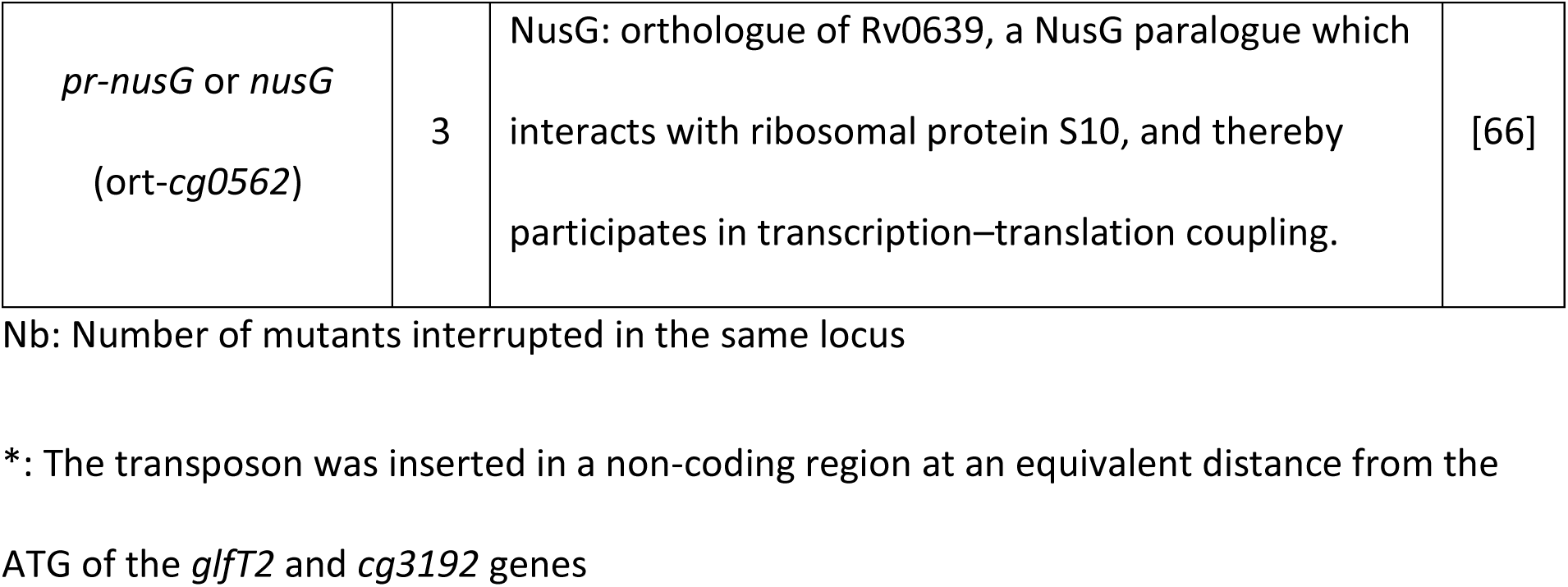
Genes interrupted by the transposon which encode characterized functions.

These results unambiguously showed that our immunological screen is a powerful tool for the identification of proteins involved in cell wall compound biosynthesis and, more widely, in cell wall biogenesis (Fig 3C).

### Identification of 22 putative new players in cell wall biogenesis of *C. glutamicum*

Twenty-two loci (corresponding to 30 different mutants) of unknown, or poorly characterized functions have been identified by our screening method. As shown in Table 3, approximately 60 % of the uncharacterized proteins fall into the category “function unknown” according to the EggNOG functional classification [32]. Although we have very limited (or no) indications of their function, as expected from the panel of known genes identified in Table 2, at least one half of these unknown proteins could be involved in cell wall biogenesis. To support this hypothesis, an analysis of the translated sequences showed that, while 5 correspond to hypothetical cytosolic proteins, 13 correspond to hypothetical membrane proteins and 3 to putative secreted proteins (Table 3 and Fig 3D). Thus, 73% of the unknown proteins found in this study are predicted to localize in the bacterial cell envelope (inner membrane or cell wall), a result that reinforces the idea that most of the proteins targeted by our screen are associated with cell envelope functions. It should be noted that for 2 mutants (5267 and 3464 see S2 Table), we do not know if only one gene was affected by the transposon and, if so, which one. In mutant 5267, the transposon insertion was localized both at the very beginning of the coding sequence of ort-*cg1137* but also presumably in the ort-*cg1136* promoter. Both genes encode unknown proteins. In mutant 3464, the transposon was inserted in an intergenic region at an equal distance from each of the ATG codons of ort-*cg3191* (*glfT2*) and ort-*cg3192* (encoding an unknown function). Because of its role in AG biosynthesis, it is tempting to favor an impact on *glfT2* (Table 2), but an impact on *ort*-*cg3192* cannot be excluded.

**Table 3:**
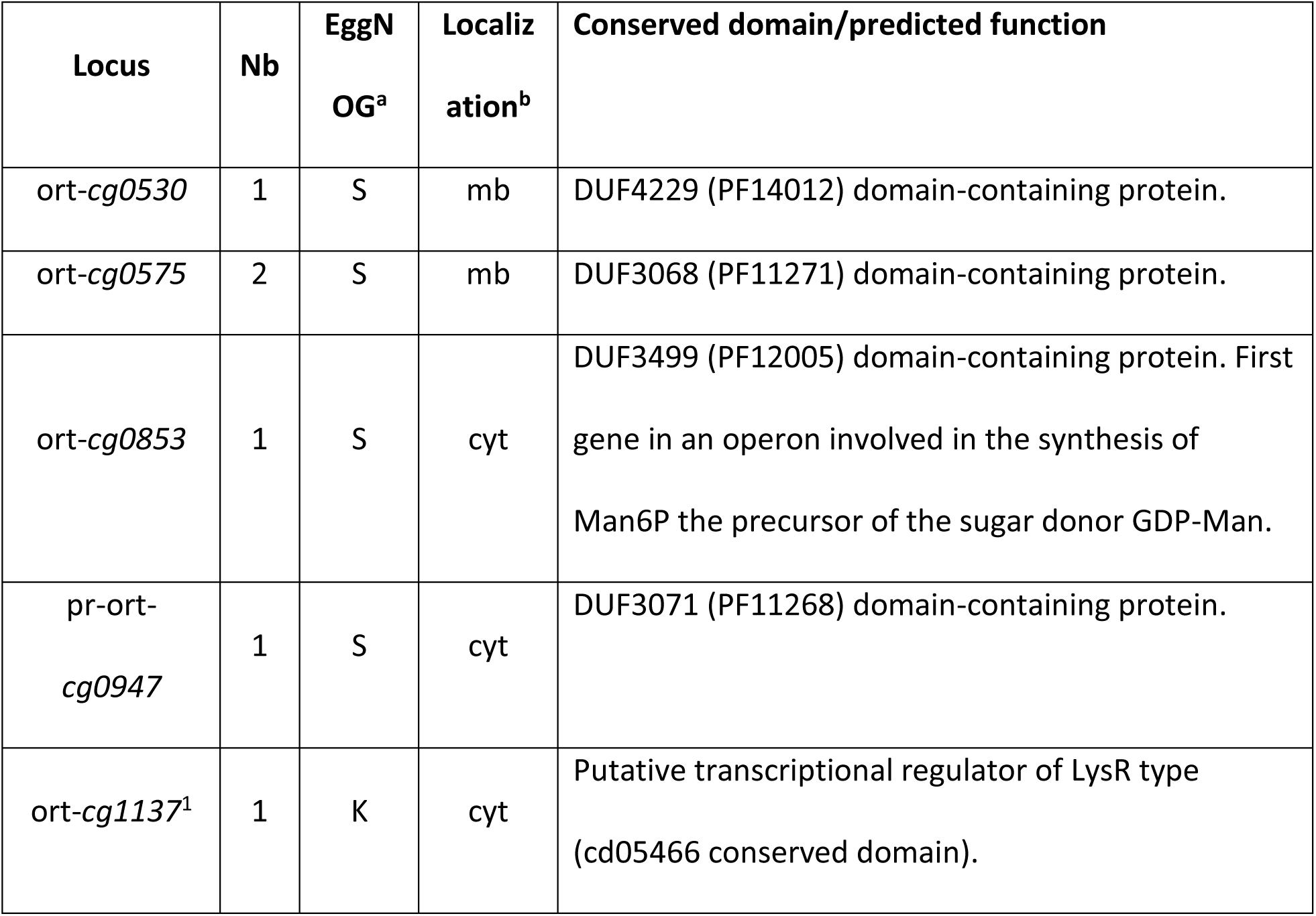
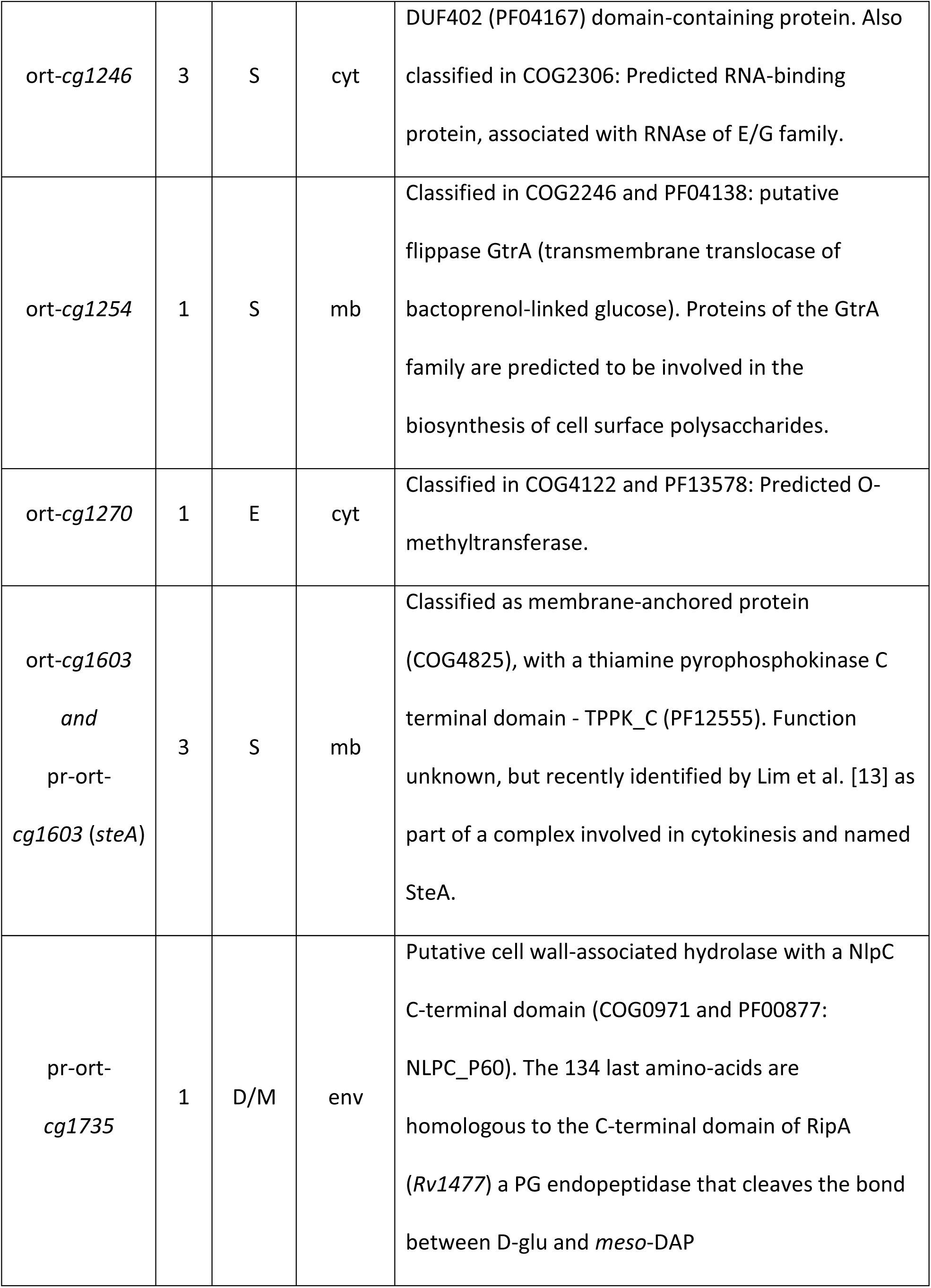
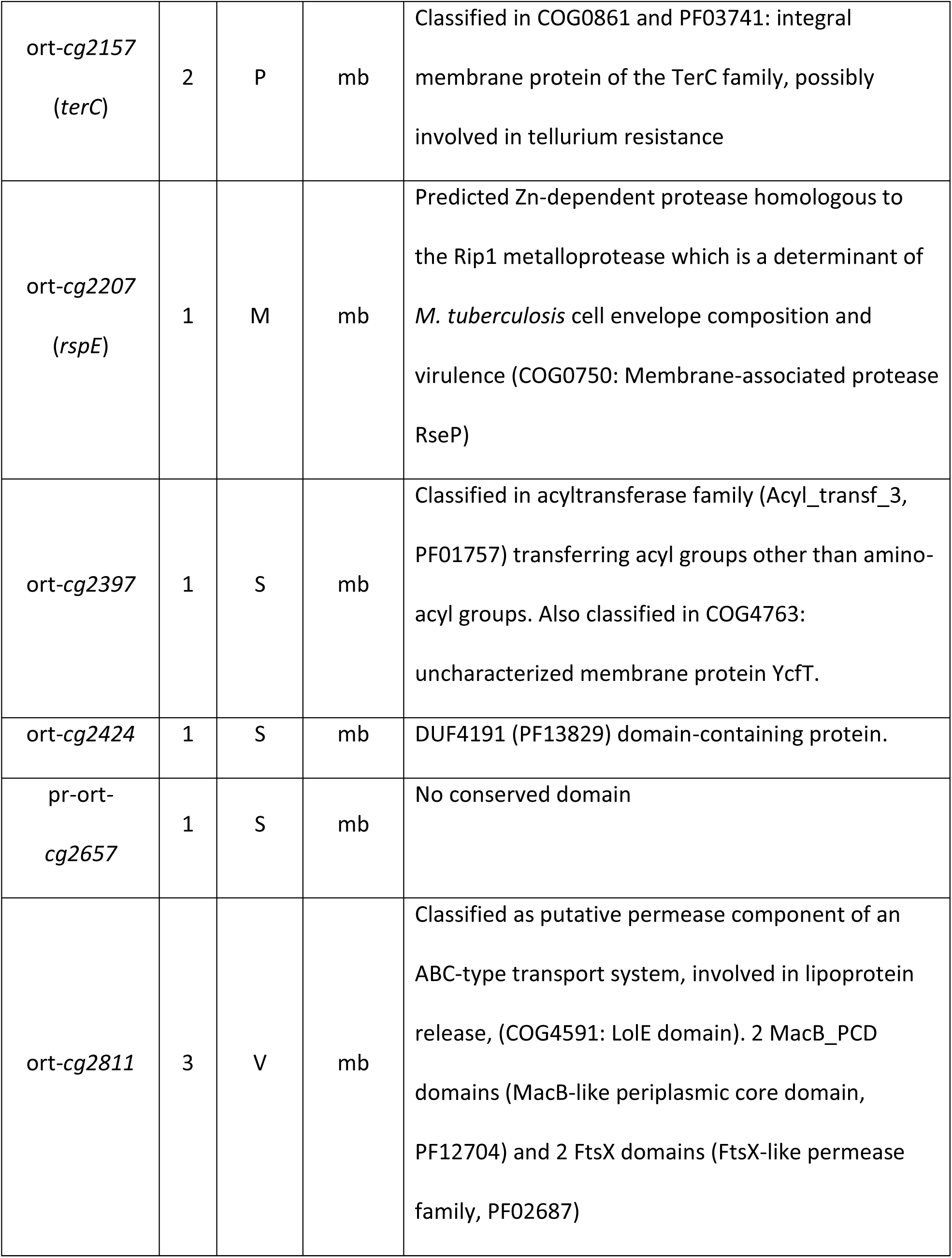
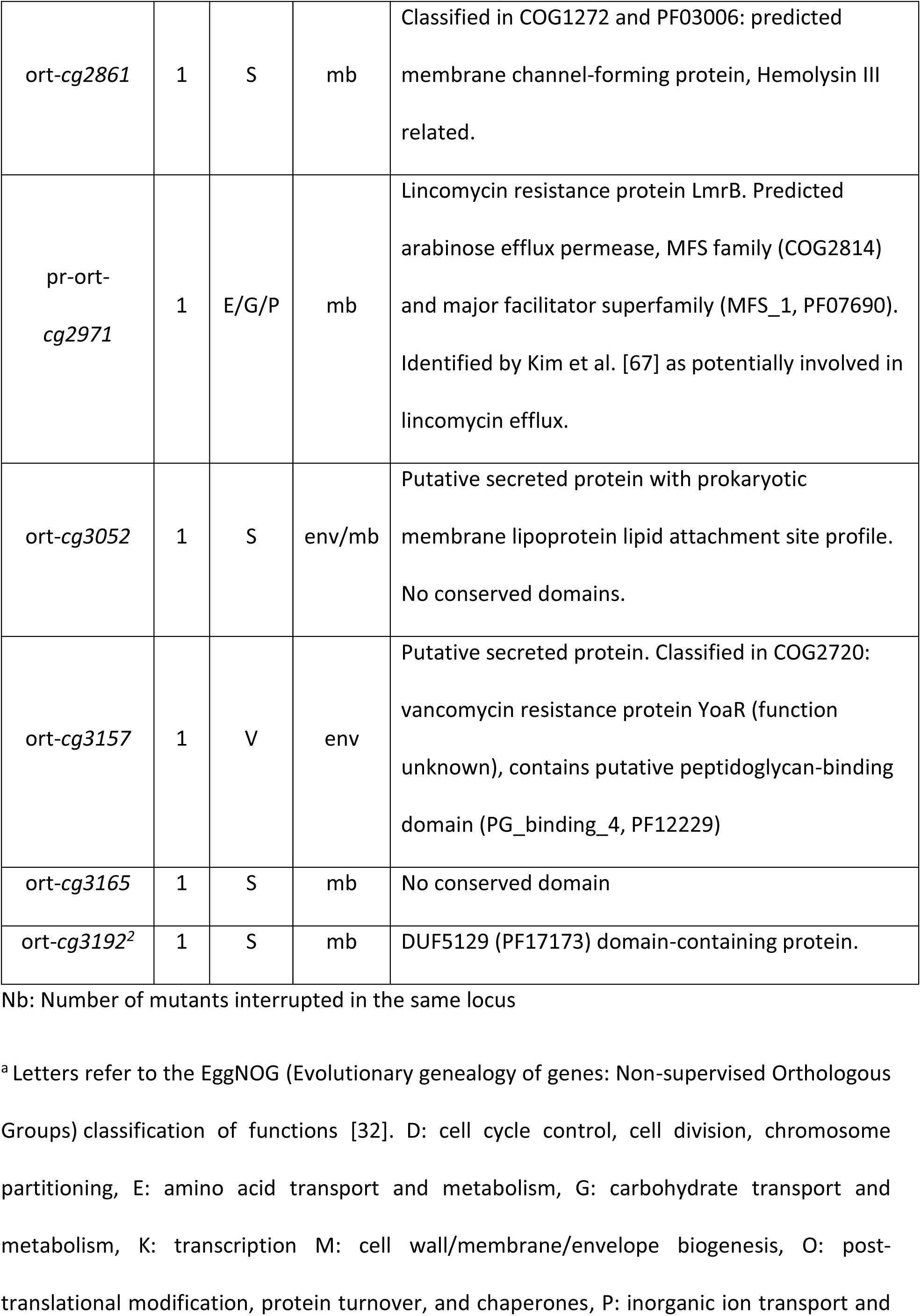
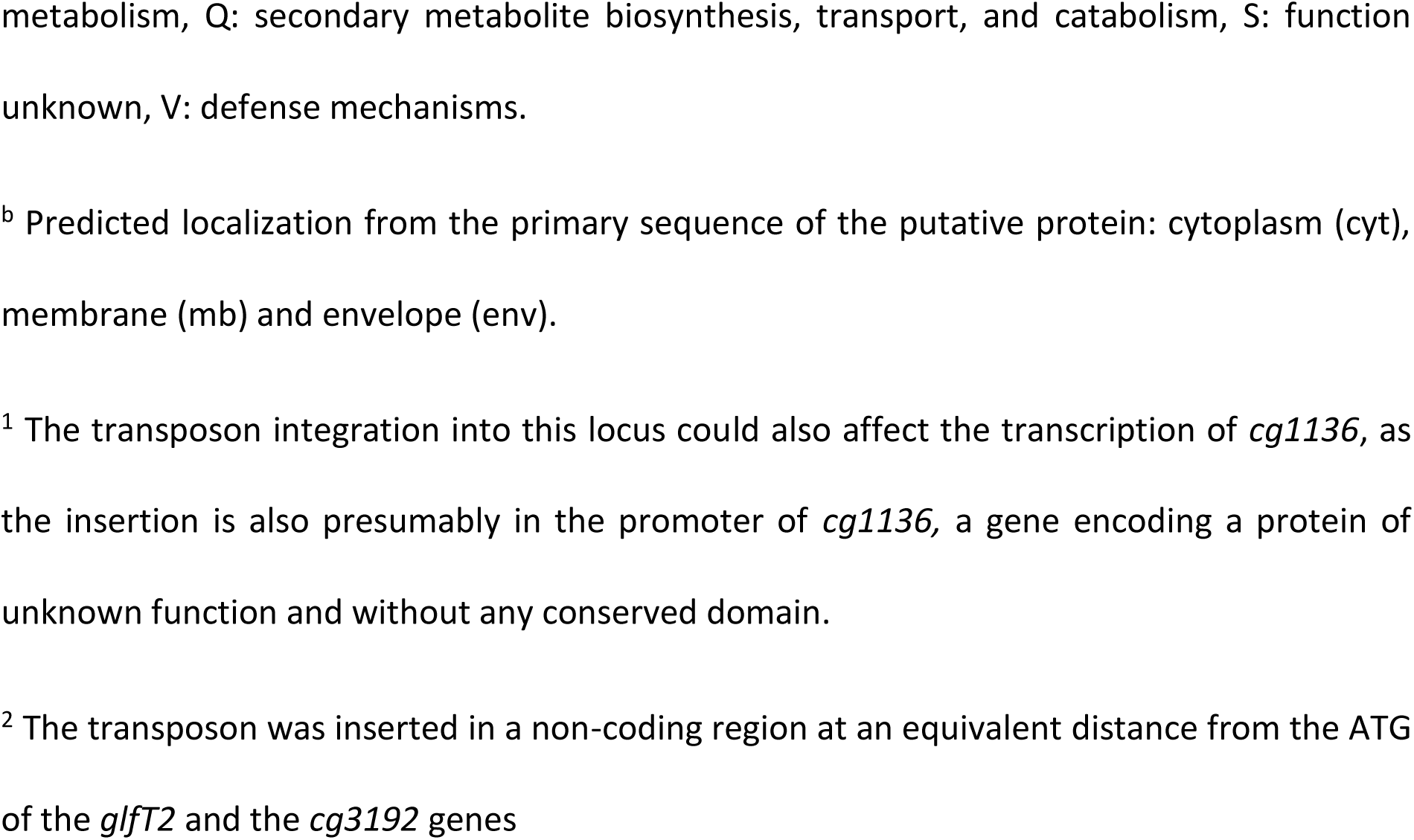
Genes interrupted by the transposon that encode poorly characterized or unknown functions.

#### Hypothetical membrane proteins

Among the 13 hypothetical membrane proteins uncovered here, 2 are most likely related to envelope dynamics. The first one is ort-Cg1603 (whose locus was inserted 3 times by the transposon) a membrane-anchored protein with a putative cytoplasmic domain and a poorly conserved thiamine diphosphokinase domain (PF12555) at its C-terminal. In ATCC13032, *cg1603* is predicted to be transcribed with *cg1604*, which encodes a protein orthologous to the mycobacterial outer membrane protein MctB, involved in copper efflux [68]. Surprisingly, *cg1603* and *cg1604* were very recently identified as conferring ethambutol hypersensitivity and cell separation defects when inactivated [13]. The authors proposed that Cg1603 (named SteA) and Cg1604 (named SteB) both localized in the inner membrane at the division site where they form a complex. This complex was hypothesized to connect other division proteins and in particular a putative periplasmic PG endopeptidase (Cg1735) that we also identified in this study. The second protein is ort-Cg2207, a putative membrane-embedded Zn-dependent protease of the RseP (Regulator of Sigma E protease) family that may play a role in cell biogenesis regulation. Indeed, the orthologue in *M. tuberculosis* is the Rip1 protein which controls 4 extracytoplasmic function (ECF) sigma (σ) factor pathways (K,L,M and D) [69, 70] and influences the lipid composition of the mycobacterial envelope, including MAs, by controlling the transcription of many specific genes [71]. It is tempting to hypothesize that Cg2207 could act on the σ^D^ pathway that controls the integrity of the cell envelope in *C. glutamicum* [72, 73]. However, it is important to note that (i) unlike SigD (the sigma factor), the anti-sigma factor RsdA is not conserved between *C. glutamicum* and *M. tuberculosis*, (ii) the σ^D^ regulons are different between the two genera [72–74] and (iii) there is no experimental evidence for a proteolytic function of Cg2207 in *vivo*.

Four loci hit by the transposon encode potential membrane transporters (ort-Cg1254, ort-Cg2811, ort-Cg2157 and ort-Cg2971). Cg1254 is annotated as a putative flippase of the GtrA family (COG2246), a group of proteins predicted to be involved in the biosynthesis or translocation of precursors of cell surface polysaccharides. Cg2811 (whose gene was inserted 3 times) is annotated as a predicted ABC-type permease transport system member, with conserved domains that belong to protein families involved in lipoprotein or lipid transport across the envelope (COG4591). In ATCC13032, *cg2811* is predicted to be transcribed with *cg2812* that encodes the putative cognate ATPase component of the system. Cg2157 (whose gene was inserted 2 times) is annotated as TerC (for Tellurite resistance protein, COG0861) and classified in the general category “inorganic ion transport and metabolism”. It is interesting to note that *cg2157* is located just downstream of the *ftsK* gene (*cg2158*) encoding a DNA translocase essential for cell division, that was also a target of the transposon (see Table 2). It appears that, neither the nature of the substrates transported by these three proteins, nor their role in the biosynthesis of the cell envelope can be deduced from these annotations. The fourth putative transporter (ort-Cg2971) is annotated as LmrB in ATCC13032 strain and was previously described to be involved in the proton-dependent efflux of the antibiotic lincomycin [67]. It is not currently known if Cg2971 transports other substrates in connection with envelope biosynthesis. Two other hypothetical membrane proteins identified here have sequences that matched with conserved domains or families (ort-Cg2397 and ort-Cg2861). Cg2397 is an uncharacterized membrane protein possessing a predicted acyl transferase domain transferring acyl groups other than amino-acyl groups (PF01757). Interestingly in ATCC13032, *cg2397* is located in the vicinity of genes encoding proteins involved in envelope biogenesis: MptA (*cg2385*), MptD (*cg2390*) MptC (*cg2393*) and PimB’ (*cg2400*) (lipoglycan biosynthesis), MytF (*cg2394*) (mycolate biosynthesis), PlsC (*cg2398*) (phospholipid biosynthesis) and Cg2401 a putative secreted PG lytic protein. Cg2861 is a predicted membrane channel-forming protein belonging to the hemolysin III family (COG1272). Finally, 5 putative membrane proteins have been identified whose sequences matched with domains of unknown function (DUF) indexed in the PFAM database (ort-Cg0530, ort-Cg0575 (2 mutants) and ort-Cg2424) or has no conserved domain (ort-Cg2657 and ort-Cg3165). Although lacking any information on their putative function, 2 of these genes were found to be located in clusters involved in the biosynthesis of cell envelope compounds implying that they are relevant in this context. Indeed, *cg0530* is surrounded by genes encoding proteins responsible for the biosynthesis of respiratory chain components (quinone, cytochrome c, heme) and *cg3165* is within a large cluster dedicated to cell envelope biosynthesis. It is interesting to note that, eight genes from this locus were inserted by the transposon (*ort*-*cg3157*, *aftD* (*ort*-*cg3161*), *mtrP* (*ort*-*cg3165*), *ort*-*cg3168*, *cg*-*pks* (*ort*-*cg3178*), *mytA* (*ort*-*cg3182*), *aftB* (*ort*-*cg3187*) and potentially *glfT2* (*ort*-*cg3191*) and/or *ort*-*cg3192* (Fig 3B).

#### Hypothetical secreted proteins

Two proteins identified in this work possess a putative sec-type signal sequence (ort-Cg1735 and ort-Cg3157) and are likely related to PG metabolism. Indeed, Cg1735 possesses a C-terminal NlpC/P60 domain generally associated with a cell wall peptidase activity [75]. Although the protein was named RipC by Lim et al. [13], it is not orthologous to the protein of the same name in *M. tuberculosis*. Nevertheless, in accordance with a putative function in PG hydrolysis, and despite the absence of enzymatic data, two studies have shown that inactivation of the protein lead to important defects in cell separation, a result that links the protein to the division process in *C. glutamicum* [13, 76]. Cg3157 possesses both a PG-binding domain (PF12229) and a VanW domain (COG2720, associated to vancomycin resistance, a PG modification), which suggests a role of Cg3157 in PG metabolism.

One protein possesses a predicted lipoprotein lipid attachment site (ort-Cg3052) but has no discernable conserved domain.

#### Hypothetical soluble non-secreted proteins

Five proteins are predicted to be non-membranous and non-secreted (ort-Cg0853, ort-Cg0947, ort-Cg1137, ort-Cg1246 and ort-Cg1270). One of them (Cg1246) could be linked to cell envelope biosynthesis. Indeed, although the putative protein has unknown function, the corresponding gene (inserted by the transposon 3 times) is part of the σ^D^ regulon that regulates mycomembrane biosynthesis and PG structure [72, 73]. It is also possible that Cg0853 is indirectly involved in the construction of the cell envelope. Indeed, in ATCC13032, its gene is surrounded by genes involved in GDP-mannose biosynthesis (*manA*, encoding a mannose-6-phosphate isomerase and *pmmA*, encoding a phosphomannomutase which form an operon with *cg0853* and *rmlA2* encoding a mannose-1-phosphate guanylyl transferase). As this nucleoside-diphosphate-sugar is an important provider of mannose for lipopolysaccharide and protein mannosylation in the cell envelope, disruption of its synthesis could have negative effects on cell wall integrity. There are no current indications linking the three remaining proteins (ort-Cg1270, ort-Cg1137 and ort-Cg0947) to any process related to the cell envelope: Cg1270 is annotated as a putative O-methyltransferase (COG4122), Cg1137 as a putative regulator of the LysR family and Cg0947 a protein with a conserved domain of unknown function.

### Approximately half of the non-characterized proteins are conserved among the *Corynebacteriales*

Since *Corynebacteriales* share unique properties in relation to their cell wall, we determined whether the proteins of unknown function identified in this work are conserved within this order. We chose 5 species, representative of the main genera that compose this order (*M. tuberculosis*, *Rhodococcus erythropolis*, *Nocardia farcinica*, *Gordonia bronchialis*, *Tsukamurella paurometabola*) and *M. leprae* because of its highly degenerate genome and searched for orthologues among these different species using BLASTp. As shown in S3 Table, 7 proteins identified by our screen (Cg0853, Cg0947, Cg1270, Cg1603, Cg2207, Cg2424 and Cg3165) have orthologues in all 6 species, with Cg1603 and Cg2207 being the best conserved between them (score > 200). Four proteins possess orthologues in all species except *M. Leprae* (Cg1246, Cg2157, Cg2861 and Cg2971) with Cg2157 being the best conserved (score > 200). Thus, about half of the proteins found in this study is conserved among *Corynebacteriales* and in particular in *M. tuberculosis*. Of the 12 *M. tuberculosis* orthologous proteins, 5 are essential according to Sasseti et al. (Rv0883c, Rv1697, Rv2219, Rv0226c, Rv0224c) [77]. Five proteins appear to be specific to the *Corynebacterium* genera (Cg0575, Cg1254, Cg2397, Cg3052 and Cg3192) with Cg3052 restricted to C. *glutamicum*.

### Cg1246 is very likely involved in mycolic acid metabolism

To identify transposon insertion in genes that are potentially involved in mycolate biosynthesis, we performed TLC analysis of organic solvent-extractable lipids from the 30 mutants interrupted in loci encoding proteins of unknown function and searched for those with altered mycolate profiles (data not shown). Three mutants (4954, 6935 and 7968, S2 Table) clearly showed a significant decrease in TMM (Fig 4A). The 3 mutants were all interrupted in a single gene, ort-*cg1246*. We thus attempted to delete *cg1246* in the *C. glutamicum* RES167 strain. For that purpose, a non-replicative pK18mobsac-derivative plasmid was constructed (pK18mobsacΔ1246) carrying sequences adjacent to the gene [24]. Kanamycin-sensitive and sucrose-resistant clones resulting from two recombination events between the plasmid and chromosome were easily obtained. Colonies in which the second recombination event led to proper deletion of the desired DNA fragment were identified using PCR with primers designed up- and downstream of this fragment. One mutant, corresponding to the expected deletion, was chosen for further characterization (Δ1246 strain). As expected from the results obtained with the ort-*cg1246* interrupted mutants, the TLC profile of the Δ1246 extractable lipids showed a significant decrease in the TMM pool as compared to the WT strain, which was restored by complementation with a plasmid bearing a copy of the wild type *cg1246* gene (Fig 4B). Quantification of trehalose lipids, performed after radiolabelling with 1-^14^C palmitate, and TLC analysis of the extractable lipids, confirmed the importance of the TMM deficiency in the Δ1246 strain, which could be estimated as four-fold when compared to the WT strain in exponentially growing cells (Fig 4C). In contrast, the TDM pool remained comparable between these two strains under the same growth conditions. Thus, the Δ1246 mutant displayed a TMM/TDM ratio lower than that of the WT cells in exponential growth (Fig 4D). In stationary phase, due to the very low level of TMM naturally present in the cell wall, the effects of *cg1246* inactivation were much less visible and led to a TMM/TDM ratio nearly similar to that of the wild type (data not shown). From these data, we conclude that Cg1246 has an important impact on the pool of TMM. How Cg1246 is connected to the mycolate pool could not be deduced from these preliminary data nor from the protein sequence itself. However, two lines of evidence argue in favor of a direct involvement of Cg1246 in mycolate biosynthesis. First, as highlighted above, the gene was found to be upregulated by σ^D^, the ECF sigma factor which was shown to regulate MA biosynthesis in *C. glutamicum* [72, 73]. Second Cg1246 is well conserved in most of *Corynebacteriales* members and is mostly associated with bacteria of this order, a specificity that is in accordance with a metabolic function (mycolate biosynthesis) that is unique to bacteria of this taxon.

**Fig 4.**
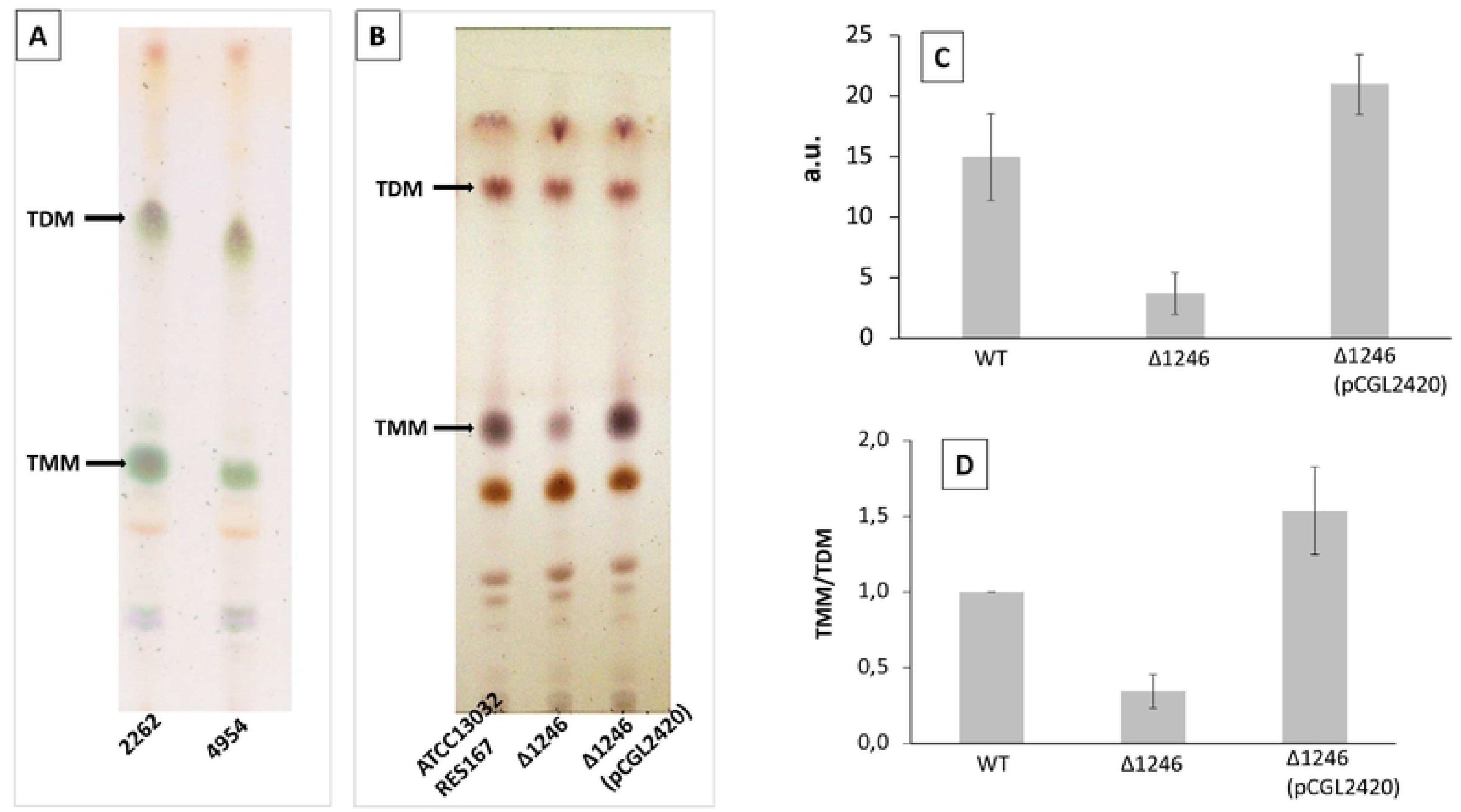
Lipid analyses of Cg1246 mutants. (A) and (B): TLC analysis of crude lipid extracts isolated from exponentially growing *cg1246*-inactivated mutants. Lipids were extracted as described in Materials and Methods and comparable amounts were loaded on TLC plates. (A): parental strain *C. glutamicum* 2262 and its isogenic mutant 4954. Glycolipid spots were visualized by spraying 0.2 % anthrone in H_2_SO_4_, followed by charring. (B): parental strain ATCC13032 and its isogenic mutant strains Δ1246 and Δ1246(pCGL2420). Lipids were visualized after immersion of the plate in 10% H_2_SO_4_ in ethanol, followed by charring. Arrows indicate the position of trehalose mono- and di-mycolate (TMM and TDM), respectively. (C) and (D): lipid quantification of RES167 strain (WT) and its derivative Δ1246 and Δ1246(pCGL2420). Exponentially growing cells were labeled with [1-^14^C]palmitate for 1.5 h and then extracted with organic solvents as described in Materials and Methods and, the radiolabeled extractable lipids were analyzed by TLC-phosphorImaging. (C): TMM quantities in arbitrary units (a.u.). (D): TMM/TDM. The relative radioactivity incorporated into TMM and TDM was determined for each strain, the ratio was calculated and normalized to 1 for the WT strain (RES167). The values in (C) and (D) are the means ± SD of at least three independent experiments.

## Conclusions

Compared to the large knowledge base acquired concerning cell envelope biogenesis in Gram-negative bacteria, little is known about the biosynthesis and assembly of the didermic cell envelope of *Corynebacteriales*. However, since the direct visualization of a mycolate containing outer membrane [5, 6], an ever-increasing number of studies have been published on the biosynthesis and assembly of the cell envelope of these bacteria. An important part of the current knowledge on this subject came from the use of *C. glutamicum* because, unlike mycobacteria, most of the known genes involved in MA and AG biosynthesis are not essential in this bacterium [3]. It is thus natural that *C. glutamicum* constitutes a model of choice to search for new genes involved in envelope biosynthesis processes. In this context, we used a classical transposon mutagenesis strategy, combined with an original immunological screening. We retained 80 mutants out of approximately 10,000 screened colonies, which corresponded to 55 independent loci. The effectiveness of our screening method was attested to by the identification of 34 interrupted loci encoding already known functions, more than half of which is involved in cell envelope biogenesis, *i.e*. biosynthesis of cell envelope components as well as cell envelope dynamics and assembly including cell division. It is therefore legitimate to assume that a significant part of the 22 loci of unknown or poorly characterized function that we identified in this work are also involved in cell envelope biogenesis. Consistent with this hypothesis is the fact that some of the genes identified in this study were also found by Lim et al. [13] in their large-scale transposon mutagenesis study associated with sensitivity to ethambutol (*ste* genes). Indeed, among the 49 *ste* genes, 12 were also found here (indicated in S2 Table), five of which are uncharacterized genes (cg0575, cg0853, cg1254, cg2811, cg3165) and two encode poorly characterized proteins linked to cell division (cg1603 and cg1735). Five additional *ste* mutations were found in predicted operons that were also inactivated by our transposon and detected by our screen (also indicated in S2 Table).

Here we identified *cg1246*, a gene encoding a protein of unknown function well conserved in *Corynebacteriales* and relatively specific to this order. We characterized the corresponding mutant and showed the involvement of Cg1246 in MA metabolism. The functional characterization of this protein is currently underway.

## Acknowledgments

We are very grateful to Pr N. Bayan and Dr M. Daffé for helpful discussions. We want to acknowledge Pr M. Dubow for critical reading of the manuscript and corrections. We thanks M. Millot, L. Graffagnino and C. Couteau for all their technical help.

## Supporting information

**S1 Fig. Examples of membranes obtained after immunological screening.** (A): Example of a membrane obtained after a first round of immunological screening. Nighty-six colonies corresponding to 96 different mutant strains, were grown on a nitrocellulose membrane layered on a BHI-plate. The membrane was treated with anti-AG antiserum and revealed as described in Materials and Methods. The lower line corresponds to control strains: 2 colonies of the WT strain 2262 and 3 colonies representing positive controls (Cg-Pks^-^ and MytA^-^, strains inactivated in *cg-pks* and *mytA* genes respectively). The white arrows indicate 3 colonies selected for a second round of screening. (B): Example of a second-round immunological screening with mutants selected from the first round of immunological screening. On the top: the nitrocellulose membrane on which colonies have grown, on the bottom: the nitrocellulose membrane corresponding to the imprint of the agar plate. For more readability, the “imprint” sheet has been flipped to be read in the same direction as the sheet on which mutants grew. White and black arrows indicate mutants to which a score of 2 or 1 have been assigned, respectively.

**S2 Fig. SDS-PAGE analysis of cell wall and extracellular proteins from the strain *C. glutamicum* 2262 (WT) and four different mutant strains**. Procedures were performed as described in Materials and Methods. In this example, the mutants whose numbers are written in red (strains n° 731, 807 and 3230) have been scored 1 because of a visible alteration of their cell wall and extracellular protein profiles. The mutant 3137 was scored 0 because of the similarity of its protein profiles with those of the WT strain. S, supernatant containing the secreted proteins; CW, cell-wall fraction; MM, molecular mass markers (in kDa).

**S1 Table: Primers used in this study**

**S2 Table: List of mutant strains and genes interrupted by the transposon**. Transposon insertion sites and locus tags are given in relation to the genome of the SCgG2 strain (NC_021352). Gene in operon are predicted from the transcriptome study of *C. glutamicum* ATCC13032 published by Pfeifer-sancar et al. [33]. *ste* hits correspond to genes identified by Lim et al. [13] from a screen based on an increased sensitivity to ethambutol of a library of mutants. Scores were assigned as described in the text: 1 or 2 for the immunological signal (see Fig. S1), 1 or 0 for the cell wall and secreted protein profiles (see Fig. S2) and 1 or 0 if the mutant exhibited at least one phenotypical or growth particularity (see text).

TSS: transcription start sites, PO: primary operon, HP: hypothetical protein, HMP: hypothetical membrane protein, TMS: transmembrane segment, aa: amino acids. ND: not determined

**S3 Table: List of *Corynebacteriales* orthologues of the uncharacterized proteins found in this study.** Search for orthologous proteins in *Corynebacteriales* genomes was performed using BLASTp online software at the NCBI. Six different species were chosen for this analysis: *Mycobacterium tuberculosis* H37Rv (NCBI:txid83332), *Mycobacterium leprae* TN (NCBI:txid272631), *Rhodococcus erythropolis* SK121 (NCBI:txid596309), *Nocardia farcinica* IFM 10152 (NCBI:txid247156), *Gordonia bronchialis* DMS 43247 (NCBI:txid526226), *Tsukamurella paurometabola* DMS 20162 (NCBI:txid521096). The highest alignment score (Max score), the percentage of sequence covered by alignment (Query cover), the Expect value (E value) and the percent identity between the sequences (Per. ident) are obtained from the BLAST result page, given by the online software at https://blast.ncbi.nlm.nih.gov.

## References

1. Goodfellow M, Jones AL. Corynebacteriales ord. nov. Bergey’s Manual of Systematics of Archaea and Bacteria. Chichester, UK: John Wiley & Sons, Ltd; 2015. pp. 1–14. doi:10.1002/9781118960608.obm00009

2. Daffé M, Marrakchi H. Unraveling the Structure of the Mycobacterial Envelope. Microbiol Spectr. 2019;7. doi:10.1128/microbiolspec.GPP3-0027-2018

3. Houssin C, de Sousa d’Auria C, Constantinesco F, Dietrich C, Labarre C, Bayan N. Architecture and Biogenesis of the Cell Envelope of Corynebacterium glutamicum. In: Inui M, Toyoda K, editors. Corynebacterium glutamicum: Biology and Biotechnology. Cham: Springer International Publishing; 2020. pp. 25–60. doi:10.1007/978-3-030-39267-3_2

4. Marrakchi H, Lanéelle M-A, Daffé M. Mycolic acids: structures, biosynthesis, and beyond. Chem Biol. 2014;21: 67–85. doi:10.1016/j.chembiol.2013.11.011

5. Zuber B, Chami M, Houssin C, Dubochet J, Griffiths G, Daffé M. Direct visualization of the outer membrane of mycobacteria and corynebacteria in their native state. J Bacteriol. 2008;190: 5672–80. doi:10.1128/JB.01919-07

6. Hoffmann C, Leis A, Niederweis M, Plitzko JM, Engelhardt H. Disclosure of the mycobacterial outer membrane: cryo-electron tomography and vitreous sections reveal the lipid bilayer structure. Proc Natl Acad Sci U S A. 2008;105: 3963–7. doi:10.1073/pnas.0709530105

7. Jarlier V, Nikaido H. Mycobacterial cell wall: structure and role in natural resistance to antibiotics. FEMS Microbiol Lett. 1994;123: 11–8. doi:10.1111/j.1574-6968.1994.tb07194.x

8. Bhat ZS, Rather MA, Maqbool M, Lah HU, Yousuf SK, Ahmad Z. Cell wall: A versatile fountain of drug targets in Mycobacterium tuberculosis. Biomed Pharmacother. 2017;95: 1520–1534. doi:10.1016/j.biopha.2017.09.036

9. Portevin D, De Sousa-D’Auria C, Houssin C, Grimaldi C, Chami M, Daffé M, et al. A polyketide synthase catalyzes the last condensation step of mycolic acid biosynthesis in mycobacteria and related organisms. Proc Natl Acad Sci U S A. 2004;101: 314–9. doi:10.1073/pnas.0305439101

10. Alderwick LJ, Radmacher E, Seidel M, Gande R, Hitchen PG, Morris HR, et al. Deletion of Cg-emb in *Corynebacterianeae* leads to a novel truncated cell wall arabinogalactan, whereas inactivation of Cg-ubiA results in an arabinan-deficient mutant with a cell wall galactan core. J Biol Chem. 2005;280: 32362–71. doi:10.1074/jbc.M506339200

11. Vincent AT, Nyongesa S, Morneau I, Reed MB, Tocheva EI, Veyrier FJ. The Mycobacterial Cell Envelope: A Relict From the Past or the Result of Recent Evolution? Front Microbiol. 2018;9: 2341. doi:10.3389/fmicb.2018.02341

12. Wang C, Hayes B, Vestling MM, Takayama K. Transposome mutagenesis of an integral membrane transporter in Corynebacterium matruchotii. Biochem Biophys Res Commun. 2006;340: 953–60. doi:10.1016/j.bbrc.2005.12.097

13. Lim HC, Sher JW, Rodriguez-Rivera FP, Fumeaux C, Bertozzi CR, Bernhardt TG. Identification of new components of the RipC-FtsEX cell separation pathway of Corynebacterineae. PLOS Genet. 2019;15: e1008284. doi:10.1371/journal.pgen.1008284

14. Mishra AK, Alderwick LJ, Rittmann D, Wang C, Bhatt A, Jacobs WR, et al. Identification of a novel alpha(1-->6) mannopyranosyltransferase MptB from Corynebacterium glutamicum by deletion of a conserved gene, NCgl1505, affords a lipomannan- and lipoarabinomannan-deficient mutant. Mol Microbiol. 2008;68: 1595–613. doi:10.1111/j.1365-2958.2008.06265.x

15. Puech V, Bayan N, Salim K, Leblon G, Daffé M. Characterization of the in vivo acceptors of the mycoloyl residues transferred by the corynebacterial PS1 and the related mycobacterial antigens 85. Mol Microbiol. 2000;35: 1026–41. doi:10.1046/j.1365-2958.2000.01738.x

16. De Sousa-D’Auria C, Kacem R, Puech V, Tropis M, Leblon G, Houssin C, et al. New insights into the biogenesis of the cell envelope of corynebacteria: identification and functional characterization of five new mycoloyltransferase genes in *Corynebacterium glutamicum*. FEMS Microbiol Lett. 2003;224: 35–44. doi:10.1016/S0378-1097(03)00396-3

17. Bonamy C, Guyonvarch A, Reyes O, David F, Leblon G. Interspecies electro-transformation in Corynebacteria. FEMS Microbiol Lett. 1990;54: 263–9. doi:10.1016/0378-1097(90)90294-z

18. Soual-Hoebeke E, De Sousa-D’Auria C, Chami M, Baucher MF, Guyonvarch A, Bayan N, et al. S-layer protein production by *Corynebacterium* strains is dependent on the carbon source. Microbiology. 1999;145: 3399–3408. doi:10.1099/00221287-145-12-3399

19. Dusch N, Pühler A, Kalinowski J. Expression of the Corynebacterium glutamicum panD gene encoding L-aspartate-alpha-decarboxylase leads to pantothenate overproduction in Escherichia coli. Appl Environ Microbiol. 1999;65: 1530–9. doi:10.1128/AEM.65.4.1530- 1539.1999

20. Bonnassie S, Oreglia J, Trautwetter A, Sicard AM. Isolation and characterization of a restriction and modification deficient mutant of Brevibacterium lactofermentum. FEMS Microbiol Lett. 1990;60: 143–6. doi:10.1016/0378-1097(90)90361-s

21. Kacem R, De Sousa-D’Auria C, Tropis M, Chami M, Gounon P, Leblon G, et al. Importance of mycoloyltransferases on the physiology of Corynebacterium glutamicum. Microbiology. 2004;150: 73–84. doi:10.1099/mic.0.26583-0

22. Portevin D, de Sousa-D’Auria C, Montrozier H, Houssin C, Stella A, Lanéelle M-A, et al. The acyl-AMP ligase FadD32 and AccD4-containing acyl-CoA carboxylase are required for the synthesis of mycolic acids and essential for mycobacterial growth: identification of the carboxylation product and determination of the acyl-CoA carboxylase componen. J Biol Chem. 2005;280: 8862–74. doi:10.1074/jbc.M408578200

23. Bonamy C, Labarre J, Cazaubon L, Jacob C, Le Bohec F, Reyes O, et al. The mobile element IS1207 of Brevibacterium lactofermentum ATCC21086: isolation and use in the construction of Tn5531, a versatile transposon for insertional mutagenesis of Corynebacterium glutamicum. J Biotechnol. 2003;104: 301–9. doi:10.1016/s0168-1656(03)00150-0

24. Schäfer A, Tauch A, Jäger W, Kalinowski J, Thierbach G, Pühler A. Small mobilizable multi-purpose cloning vectors derived from the Escherichia coli plasmids pK18 and pK19: selection of defined deletions in the chromosome of Corynebacterium glutamicum. Gene. 1994;145: 69–73. doi:10.1016/0378-1119(94)90324-7

25. Peyret JL, Bayan N, Joliff G, Gulik-Krzywicki T, Mathieu L, Schechter E, et al. Characterization of the cspB gene encoding PS2, an ordered surface-layer protein in Corynebacterium glutamicum. Mol Microbiol. 1993;9: 97–109. doi:10.1111/j.1365-2958.1993.tb01672.x

26. Ausubel F, Brent R, Kingston R, Moore D, Seidman J, Smith J, et al. Current protocols in molecular biology. New York: Wiley Interscience; 1987.

27. Daffe M, Brennan PJ, McNeil M. Predominant structural features of the cell wall arabinogalactan of Mycobacterium tuberculosis as revealed through characterization of oligoglycosyl alditol fragments by gas chromatography/mass spectrometry and by 1H and 13C NMR analyses. J Biol Chem. 1990;265: 6734–43.

28. Green MR, Sambrook J. Inverse Polymerase Chain Reaction (PCR). Cold Spring Harb Protoc. 2019;2019: pdb.prot095166. doi:10.1101/pdb.prot095166

29. Das S, Noe JC, Paik S, Kitten T. An improved arbitrary primed PCR method for rapid characterization of transposon insertion sites. J Microbiol Methods. 2005;63: 89–94. doi:10.1016/j.mimet.2005.02.011

30. Marchler-Bauer A, Bo Y, Han L, He J, Lanczycki CJ, Lu S, et al. CDD/SPARCLE: functional classification of proteins via subfamily domain architectures. Nucleic Acids Res. 2017;45: D200–D203. doi:10.1093/nar/gkw1129

31. Chen I-MA, Chu K, Palaniappan K, Pillay M, Ratner A, Huang J, et al. IMG/M v.5.0: an integrated data management and comparative analysis system for microbial genomes and microbiomes. Nucleic Acids Res. 2019;47: D666–D677. doi:10.1093/nar/gky901

32. Huerta-Cepas J, Szklarczyk D, Heller D, Hernández-Plaza A, Forslund SK, Cook H, et al. eggNOG 5.0: a hierarchical, functionally and phylogenetically annotated orthology resource based on 5090 organisms and 2502 viruses. Nucleic Acids Res. 2019;47: D309–D314. doi:10.1093/nar/gky1085

33. Pfeifer-Sancar K, Mentz A, Rückert C, Kalinowski J. Comprehensive analysis of the Corynebacterium glutamicum transcriptome using an improved RNAseq technique. BMC Genomics. 2013;14: 888. doi:10.1186/1471-2164-14-888

34. Bou Raad R, Méniche X, de Sousa-d’Auria C, Chami M, Salmeron C, Tropis M, et al. A deficiency in arabinogalactan biosynthesis affects *Corynebacterium glutamicum* mycolate outer membrane stability. J Bacteriol. 2010;192: 2691–700. doi:10.1128/JB.00009-10

35. Meniche X, de Sousa-d’Auria C, Van-der-Rest B, Bhamidi S, Huc E, Huang H, et al. Partial redundancy in the synthesis of the D-arabinose incorporated in the cell wall arabinan of *Corynebacterineae*. Microbiology. 2008;154: 2315–26. doi:10.1099/mic.0.2008/016378-0

36. Kalinowski J, Bathe B, Bartels D, Bischoff N, Bott M, Burkovski A, et al. The complete Corynebacterium glutamicum ATCC 13032 genome sequence and its impact on the production of l-aspartate-derived amino acids and vitamins. J Biotechnol. 2003;104: 5–25. doi:10.1016/S0168-1656(03)00154-8

37. Tropis M, Meniche X, Wolf A, Gebhardt H, Strelkov S, Chami M, et al. The crucial role of trehalose and structurally related oligosaccharides in the biosynthesis and transfer of mycolic acids in *Corynebacterineae*. J Biol Chem. 2005;280: 26573–85. doi:10.1074/jbc.M502104200

38. Wehrmann A, Phillipp B, Sahm H, Eggeling L. Different Modes of Diaminopimelate Synthesis and Their Role in Cell Wall Integrity: a Study withCorynebacterium glutamicum. J Bacteriol. 1998;180: 3159–3165. doi:10.1128/JB.180.12.3159-3165.1998

39. Schneider J, Peters-Wendisch P, Stansen KC, Götker S, Maximow S, Krämer R, et al. Characterization of the biotin uptake system encoded by the biotin-inducible bioYMN operon of Corynebacterium glutamicum. BMC Microbiol. 2012;12: 6. doi:10.1186/1471-2180-12-6

40. Hashimoto K, Kawasaki H, Akazawa K, Nakamura J, Asakura Y, Kudo T, et al. Changes in composition and content of mycolic acids in glutamate-overproducing Corynebacterium glutamicum. Biosci Biotechnol Biochem. 2006;70: 22–30. doi:10.1271/bbb.70.22

41. Radmacher E, Alderwick LJ, Besra GS, Brown AK, Gibson KJC, Sahm H, et al. Two functional FAS-I type fatty acid synthases in *Corynebacterium glutamicum*. Microbiology. 2005;151: 2421–7. doi:10.1099/mic.0.28012-0

42. Grover S, Alderwick LJ, Mishra AK, Krumbach K, Marienhagen J, Eggeling L, et al. Benzothiazinones mediate killing of *Corynebacterineae* by blocking decaprenyl phosphate recycling involved in cell wall biosynthesis. J Biol Chem. 2014;289: 6177–87. doi:10.1074/jbc.M113.522623

43. Gande R, Dover LG, Krumbach K, Besra GS, Sahm H, Oikawa T, et al. The two carboxylases of *Corynebacterium glutamicum* essential for fatty acid and mycolic acid synthesis. J Bacteriol. 2007;189: 5257–64. doi:10.1128/JB.00254-07

44. Chalut C, Botella L, de Sousa-D’Auria C, Houssin C, Guilhot C. The nonredundant roles of two 4’-phosphopantetheinyl transferases in vital processes of Mycobacteria. Proc Natl Acad Sci U S A. 2006;103: 8511–6. doi:10.1073/pnas.0511129103

45. Rainczuk AK, Klatt S, Yamaryo-Botté Y, Brammananth R, McConville XMJ, Coppel RL, et al. Mtrp, a putative methyltransferase in corynebacteria, is required for optimal membrane transport of trehalose mycolates. J Biol Chem. 2020;295: 6108–6119. doi:10.1074/jbc.RA119.011688

46. Lea-Smith DJ, Pyke JS, Tull D, McConville MJ, Coppel RL, Crellin PK. The reductase that catalyzes mycolic motif synthesis is required for efficient attachment of mycolic acids to arabinogalactan. J Biol Chem. 2007;282: 11000–8. doi:10.1074/jbc.M608686200

47. Alderwick LJ, Birch HL, Krumbach K, Bott M, Eggeling L, Besra GS. AftD functions as an α1 → 5 arabinofuranosyltransferase involved in the biosynthesis of the mycobacterial cell wall core. Cell Surf (Amsterdam, Netherlands). 2018;1: 2–14. doi:10.1016/j.tcsw.2017.10.001

48. Seidel M, Alderwick LJ, Birch HL, Sahm H, Eggeling L, Besra GS. Identification of a novel arabinofuranosyltransferase AftB involved in a terminal step of cell wall arabinan biosynthesis in *Corynebacterianeae*, such as *Corynebacterium glutamicum* and *Mycobacterium tuberculosis*. J Biol Chem. 2007;282: 14729–40. doi:10.1074/jbc.M700271200

49. Kremer L, Dover LG, Morehouse C, Hitchin P, Everett M, Morris HR, et al. Galactan biosynthesis in *Mycobacterium tuberculosis*. Identification of a bifunctional UDP-galactofuranosyltransferase. J Biol Chem. 2001;276: 26430–40. doi:10.1074/jbc.M102022200

50. Donovan C, Bramkamp M. Cell division in Corynebacterineae. Front Microbiol. 2014;5: 132. doi:10.3389/fmicb.2014.00132

51. Valbuena N, Letek M, Ordóñez E, Ayala J, Daniel RA, Gil JA, et al. Characterization of HMW-PBPs from the rod-shaped actinomycete *Corynebacterium glutamicum*: peptidoglycan synthesis in cells lacking actin-like cytoskeletal structures. Mol Microbiol. 2007;66: 643–57. doi:10.1111/j.1365-2958.2004.05943.x

52. Oikawa T, Tauch A, Schaffer S, Fujioka T. Expression of alr gene from Corynebacterium glutamicum ATCC 13032 in Escherichia coli and molecular characterization of the recombinant alanine racemase. J Biotechnol. 2006;125: 503–512. doi:10.1016/j.jbiotec.2006.04.002

53. Levefaudes M, Patin D, de Sousa-d’Auria C, Chami M, Blanot D, Hervé M, et al. Diaminopimelic Acid Amidation in *Corynebacteriales*: NEW INSIGHTS INTO THE ROLE OF LtsA IN PEPTIDOGLYCAN MODIFICATION. J Biol Chem. 2015;290: 13079–94. doi:10.1074/jbc.M115.642843

54. Valbuena N, Letek M, Ramos A, Ayala J, Nakunst D, Kalinowski J, et al. Morphological changes and proteome response of *Corynebacterium glutamicum* to a partial depletion of Ftsl. Microbiology. 2006;152: 2491–2503. doi:10.1099/mic.0.28773-0

55. Carrión M, Gómez MJ, Merchante-Schubert R, Dongarrá S, Ayala JA. mraW, an essential gene at the dcw cluster of Escherichia coli codes for a cytoplasmic protein with methyltransferase activity. Biochimie. 1999;81: 879–88. doi:10.1016/s0300-9084(99)00208-4

56. Huc E, de Sousa-D’Auria C, de la Sierra-Gallay IL, Salmeron C, van Tilbeurgh H, Bayan N, et al. Identification of a mycoloyl transferase selectively involved in o-acylation of polypeptides in *Corynebacteriales*. J Bacteriol. 2013;195: 4121–8. doi:10.1128/JB.00285-13

57. Dautin N, Argentini M, Mohiman N, Labarre C, Cornu D, Sago L, et al. Role of the unique, non-essential phosphatidylglycerol::prolipoprotein diacylglyceryl transferase (Lgt) in Corynebacterium glutamicum. Microbiology. 2020;166: 759–776. doi:10.1099/mic.0.000937

58. Schwinde JW, Hertz PF, Sahm H, Eikmanns BJ, Guyonvarch A. Lipoamide dehydrogenase from Corynebacterium glutamicum: molecular and physiological analysis of the lpd gene and characterization of the enzyme. Microbiology. 2001;147: 2223–31. doi:10.1099/00221287-147-8-2223

59. Caspi R, Billington R, Fulcher CA, Keseler IM, Kothari A, Krummenacker M, et al. The MetaCyc database of metabolic pathways and enzymes. Nucleic Acids Res. 2018;46: D633–D639. doi:10.1093/nar/gkx935

60. Ankri S, Serebrijski I, Reyes O, Leblon G. Mutations in the Corynebacterium glutamicum proline biosynthetic pathway: A natural bypass of the proA step. J Bacteriol. 1996;178: 4412– 4419. doi:10.1128/jb.178.15.4412-4419.1996

61. Davoudi C-F, Ramp P, Baumgart M, Bott M. Identification of Surf1 as an assembly factor of the cytochrome bc1-aa3 supercomplex of Actinobacteria. Biochim Biophys acta Bioenerg. 2019. doi:10.1016/j.bbabio.2019.06.005

62. Morosov X, Davoudi C-F, Baumgart M, Brocker M, Bott M. The copper-deprivation stimulon of Corynebacterium glutamicum comprises proteins for biogenesis of the actinobacterial cytochrome bc1-aa3 supercomplex. J Biol Chem. 2018;293: 15628–15640. doi:10.1074/jbc.RA118.004117

63. Bott M, Niebisch A. The respiratory chain of Corynebacterium glutamicum. J Biotechnol. 2003;104: 129–53. doi:10.1016/S0168-1656(03)00144-5

64. Jochmann N, Götker S, Tauch A. Positive transcriptional control of the pyridoxal phosphate biosynthesis genes pdxST by the MocR-type regulator PdxR of Corynebacterium glutamicum ATCC 13032. Microbiology. 2011;157: 77–88. doi:10.1099/mic.0.044818-0

65. Si M, Wang J, Xiao X, Guan J, Zhang Y, Ding W, et al. Ohr protects Corynebacterium glutamicum against organic hydroperoxide induced oxidative stress. PLoS One. 2015;10: 1–17. doi:10.1371/journal.pone.0131634

66. Kalyani BS, Kunamneni R, Wal M, Ranjan A, Sen R. A NusG paralogue from Mycobacterium tuberculosis, Rv0639, has evolved to interact with ribosomal protein S10 (Rv0700) but not to function as a transcription elongation-termination factor. Microbiology. 2015;161: 67–83. doi:10.1099/mic.0.083709-0

67. Kim HJ, Kim Y, Lee MS, Lee HS. Gene lmrB of Corynebacterium glutamicum confers efflux-mediated resistance to lincomycin. Mol Cells. 2001;12: 112–6.

68. Wolschendorf F, Ackart D, Shrestha TB, Hascall-Dove L, Nolan S, Lamichhane G, et al. Copper resistance is essential for virulence of Mycobacterium tuberculosis. Proc Natl Acad Sci U S A. 2011;108: 1621–6. doi:10.1073/pnas.1009261108

69. Schneider JS, Sklar JG, Glickman MS. The Rip1 protease of mycobacterium tuberculosis controls the SigD regulon. J Bacteriol. 2014;196: 2638–2645. doi:10.1128/JB.01537-14

70. Sklar JG, Makinoshima H, Schneider JS, Glickman MS. M. tuberculosis intramembrane protease Rip1 controls transcription through three anti-sigma factor substrates. Mol Microbiol. 2010;77: 605–17. doi:10.1111/j.1365-2958.2010.07232.x

71. Makinoshima H, Glickman MS. Regulation of Mycobacterium tuberculosis cell envelope composition and virulence by intramembrane proteolysis. Nature. 2005;436: 406–9. doi:10.1038/nature03713

72. Toyoda K, Inui M. Extracytoplasmic function sigma factor σD confers resistance to environmental stress by enhancing mycolate synthesis and modifying peptidoglycan structures in *Corynebacterium glutamicum*. Mol Microbiol. 2018;107: 312–329. doi:10.1111/mmi.13883

73. Taniguchi H, Busche T, Patschkowski T, Niehaus K, Pátek M, Kalinowski J, et al. Physiological roles of sigma factor SigD in *Corynebacterium glutamicum*. BMC Microbiol. 2017;17: 158. doi:10.1186/s12866-017-1067-6

74. Raman S, Hazra R, Dascher CC, Husson RN. Transcription Regulation by the Mycobacterium tuberculosis Alternative Sigma Factor SigD and Its Role in Virulence Transcription Regulation by the Mycobacterium tuberculosis Alternative Sigma Factor SigD and Its Role in Virulence †. J Bacteriol. 2004;186: 66505–6616. doi:10.1128/JB.186.19.6605

75. Anantharaman V, Aravind L. Evolutionary history, structural features and biochemical diversity of the NlpC/P60 superfamily of enzymes. Genome Biol. 2003;4: R11. doi:10.1186/gb-2003-4-2-r11

76. Tsuge Y, Ogino H, Teramoto H, Inui M, Yukawa H. Deletion of cgR_1596 and cgR_2070, encoding NlpC/P60 proteins, causes a defect in cell separation in Corynebacterium glutamicum R. J Bacteriol. 2008;190: 8204–14. doi:10.1128/JB.00752-08

77. Sassetti CM, Boyd DH, Rubin EJ. Genes required for mycobacterial growth defined by high density mutagenesis. Mol Microbiol. 2003;48: 77–84. doi:10.1046/j.1365-2958.2003.03425.x

